# Diffusion of DNA-binding species in the nucleus: A transient anomalous subdiffusion model

**DOI:** 10.1101/742171

**Authors:** M. J. Saxton

## Abstract

Single-particle tracking experiments have measured the distribution of escape times of DNA-binding species diffusing in living cells: CRISPR-Cas9, TetR, and LacI. The observed distribution is a truncated power law. One important property of this distribution is that it is inconsistent with a Gaussian distribution of binding energies. Another is that it leads to transient anomalous subdiffusion, in which diffusion is anomalous at short times and normal at long times, here only mildly anomalous. Monte Carlo simulations are used to characterize the time-dependent diffusion coefficient *D*(*t*) in terms of the anomalous exponent *α*, the crossover time *t*(*cross*), and the limits *D*(0) and *D*(∞), and to relate these quantities to the escape time distribution. The simplest interpretations identifSubdiffusion of DNA-binding speciesy the escape time as the actual binding time to DNA, or the period of 1D diffusion on DNA in the standard model combining 1D and 3D search, but a more complicated interpretation may be required. The model has several implications for cell biophysics. (*a*), The initial anomalous regime represents the search of the DNA-binding species for its target DNA sequence. (*b*), Non-target DNA sites have a significant effect on search kinetics. False positives in bioinformatic searches of the genome are potentially rate-determining in vivo. For simple binding, the search would be speeded if false-positive sequences were eliminated from the genome. (*c*), Both binding and obstruction affect diffusion. Obstruction ought to be measured directly, using as the primary probe the DNA-binding species with the binding site inactivated, and eGFP as a calibration standard among laboratories and cell types. (*d*), Overexpression of the DNA-binding species reduces anomalous subdiffusion because the deepest binding sites are occupied and unavailable. (*e*), The model provides a coarse-grained phenomenological description of diffusion of a DNA-binding species, useful in larger-scale modeling of kinetics, FCS, and FRAP.

**SIGNIFICANCE:** DNA-binding proteins such as transcription factors diffuse in the nucleus until they find their biological target and bind to it. A protein may bind to many false-positive sites before it reaches its target, and the search process is a research topic of considerable interest. Experimental results from the Dahan lab show a truncated power law distribution of escape times at these sites. We show by Monte Carlo simulations that this escape time distribution implies that the protein shows transient anomalous subdiffusion, defined as anomalous subdiffusion at short times and normal diffusion at long times. Implications of the model for experiments, controls, and interpretation of experiments are discussed.

## INTRODUCTION

Long ago I examined transient anomalous subdiffusion (TASD) due to traps (1) and the interpretation of TASD in terms of the search for the biological target (2). Experimental advances make it timely to revisit this topic. Single-particle tracking (SPT) experiments have caught up with modeling.

Three papers from the Dahan lab reported the escape times (equivalently, the binding, residence, or dwell times) for biomolecules interacting with DNA in living mammalian cells, and found a truncated power-law distribution: *(a)*, Repressor TetR in the nucleus of U2OS cells (3). A bacterial transcription factor was used with an artificially introduced bacterial target site in a mammalian cell. In general, this was an admirably careful and thorough study, with 73 pages of supplementary material (though I will still suggest more measurements that would be useful). *(b)*, Repressor LacI in nucleus of U2OS cells (4). *(c)*, CRISPR-Cas9 in the nucleus of 3T3 cells (5). Nonsense sgRNA (single-guide RNA) was used to examine the search phase rather than the bound phase. A related paper examined CRISPR-Cas9 binding to DNA curtains (6). The classic model of search by transcription factors is facilitated diffusion, in which 3D diffusion in solution alternates with 1D diffusion along the DNA (7, 8). An important result is that at the spatial resolution of the experiments, CRISPR-Cas9 used predominantly a 3D search, both in the DNA curtain and in cells (5, 6).

The change in the biological context of the model is important. My earlier work was phrased in terms of cell membranes and was done for discrete power-law distributions of trap depths on a 2D triangular lattice. The work here considers protein binding to DNA, and is done for continuous distributions of trap depths on a 3D simple cubic lattice. The changes in dimensionality, lattice, and distribution of trap depths affect the quantitative results but the qualitative picture is similar. The real difference is that the binding of transcription factors, polymerases, and repair proteins to DNA is much better characterized than the binding of membrane proteins or lipids to other membrane components.

We discuss TASD in the context of the search of a biomolecule for its biological target in the nucleus. We claim that *(a)*, the observed truncated power law distribution of escape times from traps leads to TASD on length scales much greater than the trap spacing; *(b)*, the deepest trap is the biological target of the diffusing molecule; and *(c)*, the anomalous regime represents the search for that target in a background of competing binding sites. We examine a simple diffusion model based on the observed distribution of traps, to see what the model predicts and what experiments it suggests.

We consider only diffusion here, not kinetics. Woringer and Darzacq (9) briefly reviewed the physics literature on fractal kinetics to assess its applicability to binding to DNA. Work by Slutsky et al. (10) and by Benichou et al. (11) examined anomalous diffusion in the nucleus in terms of the first passage time, so their results are directly linked to search kinetics. The question I address here is different: What do SPT measurements reveal about anomalous subdiffusion in the nucleus? Much work has been done on modeling chromatin structure, ranging from bottom-up physics starting with simple polymer models to top-down genomics starting with Hi-C results, reviewed in (12–15), for example. Examination of these models is beyond the scope of this work, except for some discussion of the general topic of obstructed diffusion.

## METHODS

Monte Carlo runs in Fortran were used to find the mean-square displacement as a function of time for tracers carrying out a random walk among TPL traps. Methods are considered in (1, 16). The Methods section of the supplement discusses these calculations in detail (Suppl. 1.1), as well as the analysis of TASD curves (Suppl. 1.2) and Monte Carlo calculations in Mathematica (Suppl. 1.3).

## RESULTS

### Analysis of transient anomalous subdiffusion (TASD)

We review the method of analyzing mean-square displacement data, either SPT or Monte Carlo, to show TASD. For the traps considered here, the effect of TASD is small when observed at the scale of a linear plot of 〈*r*^2^〉 versus time, but the structure is clearly shown in a plot of log *D*(*t*) versus log time (17). We argue that this structure is worth examining as a signature of the biological search process. In TASD, diffusion is anomalous at short times and normal at long times.

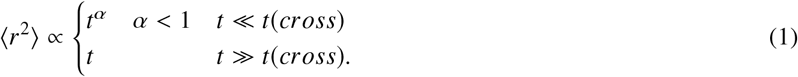

where 〈*r*^2^〉 is the mean-square displacement, *t* is time, and *t*(*cross*) is the crossover time. In other words, the diffusion coefficient *D*(*t*) = 〈*r*^2^〉/*t* is time-dependent, but reaches a constant value at large times. Normal diffusion corresponds to *α* = 1.

The simplest plot of the data is the standard plot of 〈*r*^2^〉 versus time, Fig. 1 *a*, which gives for the most part an apparently straight line of slope *D*(∞) = constant. In this example, curvature is evident for *t* ≤ 128. Taking out the asymptotic time dependence reveals the structure more clearly. A plot of 〈*r*^2^〉/*t*, Fig. 1 *b*, shows that the time range of the nonlinear regime is actually larger than Fig. 1 *a* suggests. The power-law form in Eq. 1 implies that a log-log plot is needed, and the simple plot of log 〈*r*^2^〉 versus log t in Fig. 1 *c* shows the change in slope due to the change in exponent. The plots in *(b)* and *(c)* each emphasize the initial structure, and combining them provides the clearest depiction of the structure, as shown in Fig. 1 *d*, log 〈*r*^2^〉/*t* versus log *t*. When I introduced the plots of Fig. 1 *d* (17), the point was simply to take out the asymptotic time dependence to show the initial structure, but now log 〈*r*^2^〉/*t* is generally interpreted as log *D*(*t*), suitably normalized.

**Figure 1:**
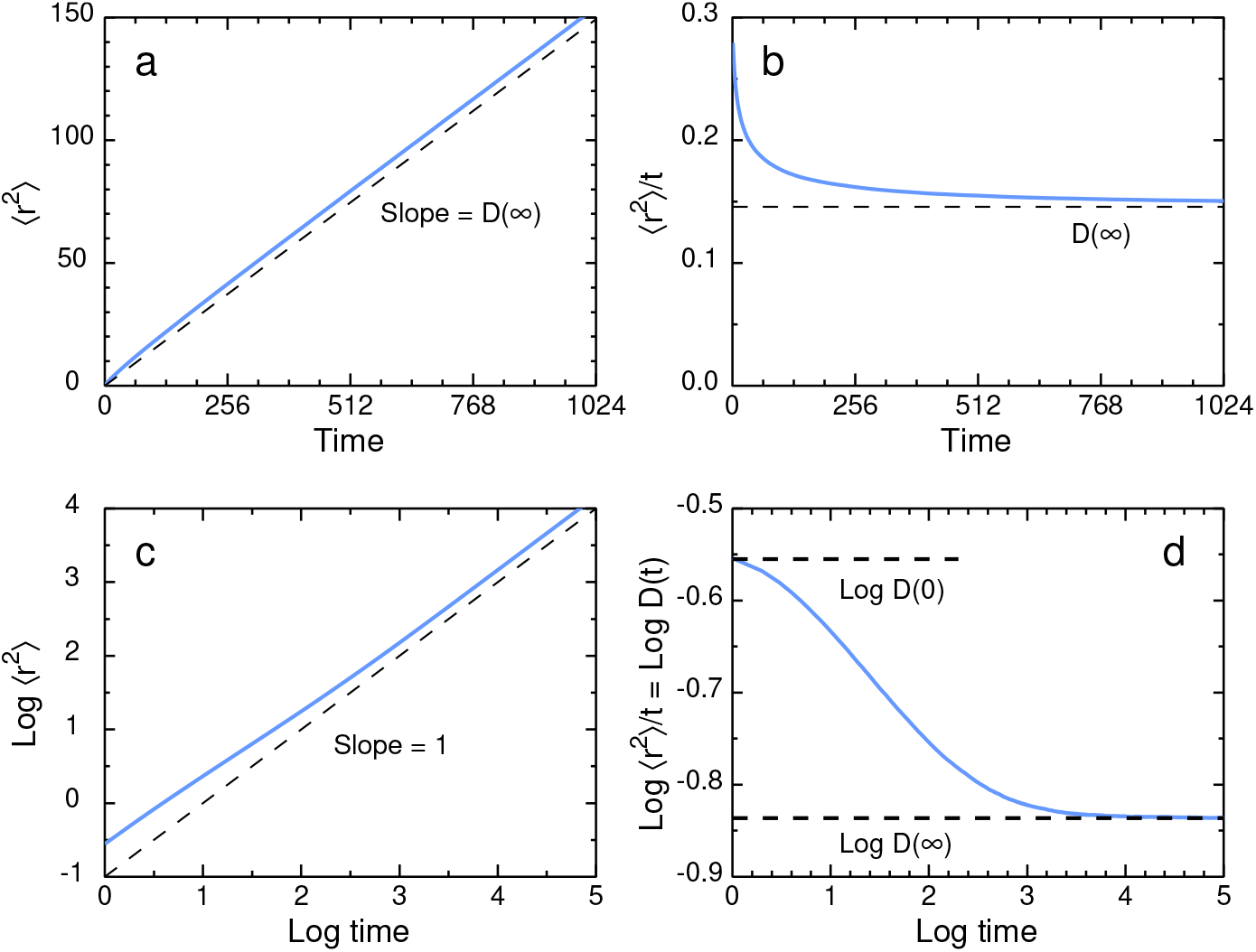
How to find the TASD structure in SPT or Monte Carlo data. Various plots of the same Monte Carlo data for a 3D random walk on a simple cubic lattice. Linear plots, *(a)*, 〈*r*^2^〉 versus *t*, and *(b)*, {*r*^2^)¦*t* versus *t*. Log-log plots, *(c)*, log 〈*r*^2^〉 versus log*t*, and *(d)*, log 〈*r*^2^〉/*t* versus log*t*. The mean-square displacement 〈*r*^2^〉 is found by ensemble averaging, with no averaging over time segments within a trajectory. This choice of averaging is essential, as discussed in the text on “aging.”

The analysis of this plot (17). is sketched in Fig. 2. The initial behavior depends on the dynamics used (Brownian versus Newtonian), and on the nature of the trap (point traps here versus multisite dead ends in a percolating cluster). The anomalous exponent *α* is given by the slope of the anomalous region around the midpoint, and the horizontal region represents normal diffusion with coefficient *D*(∞). The intersection of the slope line and the horizontal *D*(∞) line is one reasonable choice for the crossover time *t*(*cross*), as discussed in Suppl. 1.2.

**Figure 2:**
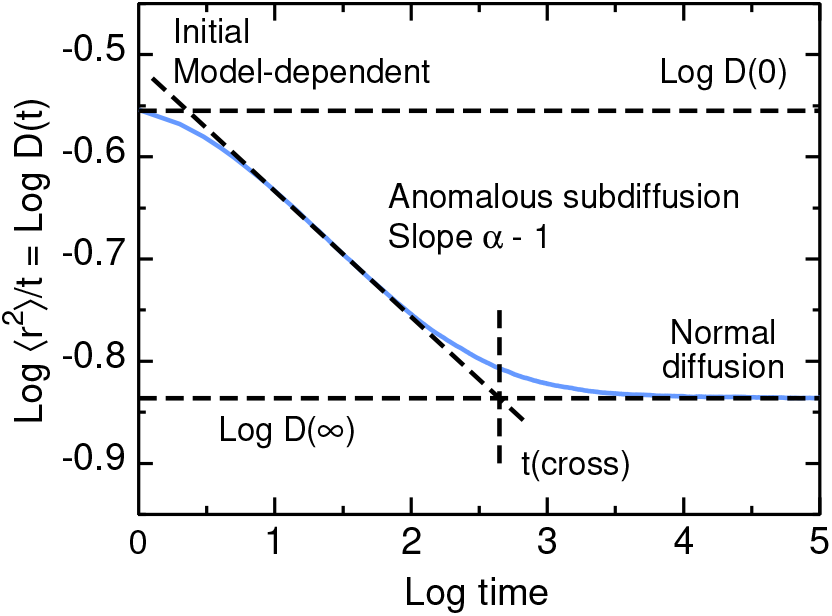
Analysis of transient anomalous subdiffusion in a plot of log 〈*r*^2^〉/*t* versus log *t*.

### Truncated power law distribution

The experimental distributions are given in terms of the survival function

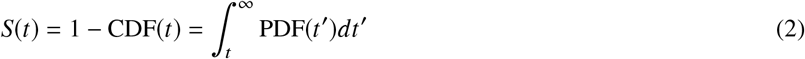

where PDF is the probability density function and CDF is the cumulative density function (18). The PDF is thus the negative derivative of the survival function, and *S*(*t*) ∝ 1/*t*^*m*-1^ yields PDF(t) ∝ 1/*t^m^*. The experimental results give a truncated power law (TPL) distribution of escape times *T*, which is defined by the probability density function

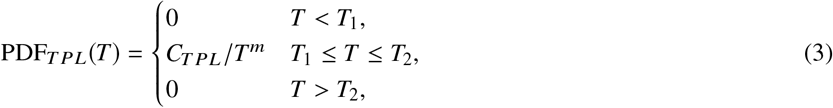

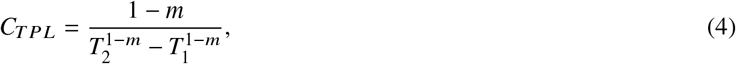

specified by a power *m* > 0, a minimum escape time *T*_1_, and a maximum *T*_2_. In this section and in Suppl. 2.2, *T* is simply a random variable, here an escape time, either the mean escape time or the observed. Power-law distributions are reviewed by Newman (19).

The power-law form is based on the experimental results of Normanno et al. (3) and the others given in the Introduction (4, 5). Parameters are given in Suppl. 2.1. The truncation at short times is somewhat subjective: How short a delay is scored as binding? Experimentalists must define a threshold to distinguish binding events from the episodes of localization that occur by chance in a pure random walk (20). The truncation at long times represents escape from a maximum trap depth. (Strictly speaking, the maximum ought to be chosen for consistency with the kinetics. If the tracer-target reaction probability is < 1, the maximum trap depth is the target depth because the tracer can escape and re-enter the target site. If the reaction probability is 1, then the maximum trap depth is the depth of the deepest false-positive site, because the trajectory ends when the tracer first encounters the target (21).)

If the PDF is truncated at *T*_1_ and *T*_2_, the CDF and survival function are also truncated there. Fig. 3 shows the shape of the distribution in linear and logarithmic plots, and Fig. 4 shows the dependence on m and on *T*_2_ as logarithmic plots. The cumulative density function of the TPL distribution is

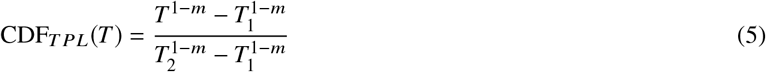

and the mean is

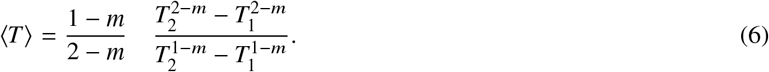

**Figure 3:**
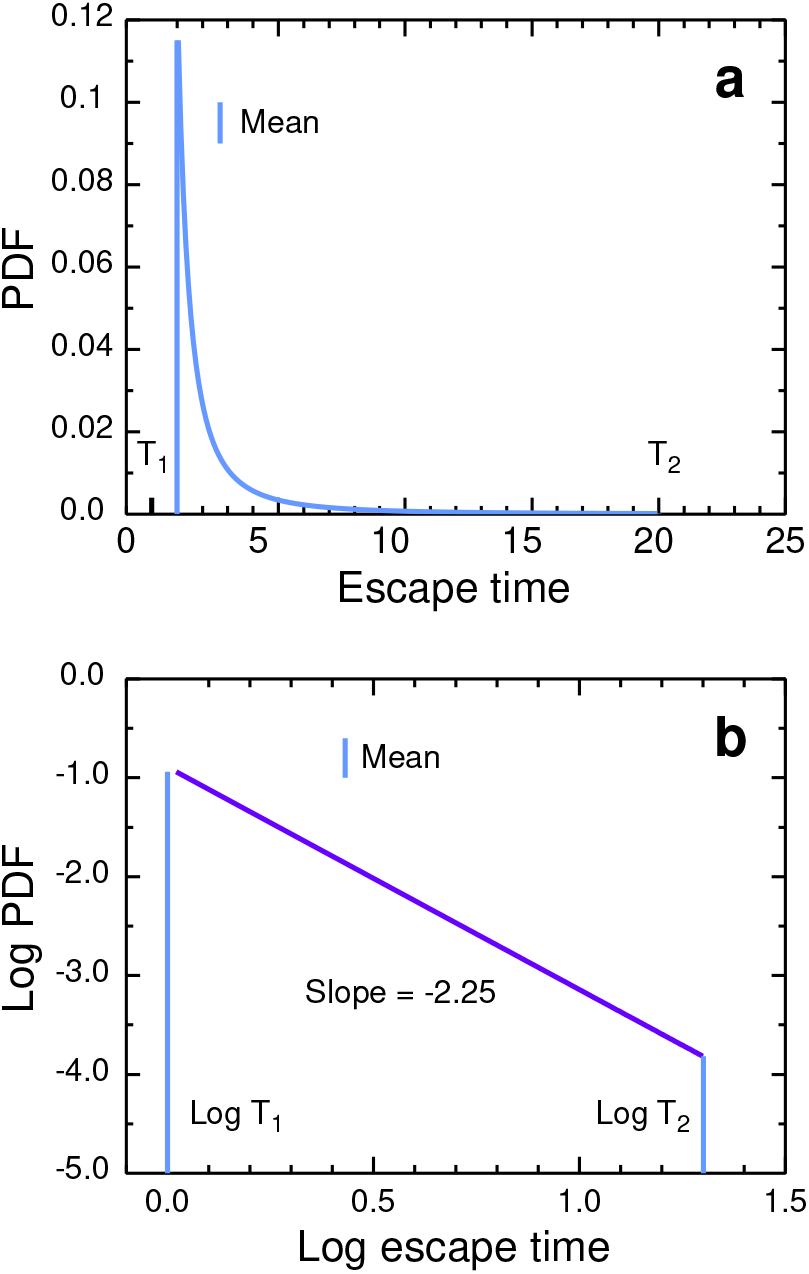
Truncated power law distribution for *m* = 2.25, *T*_1_ = 1, *T*_2_ = 200 as *(a)*, linear and *(b)*, log-log plots. Vertical lines are means.

**Figure 4:**
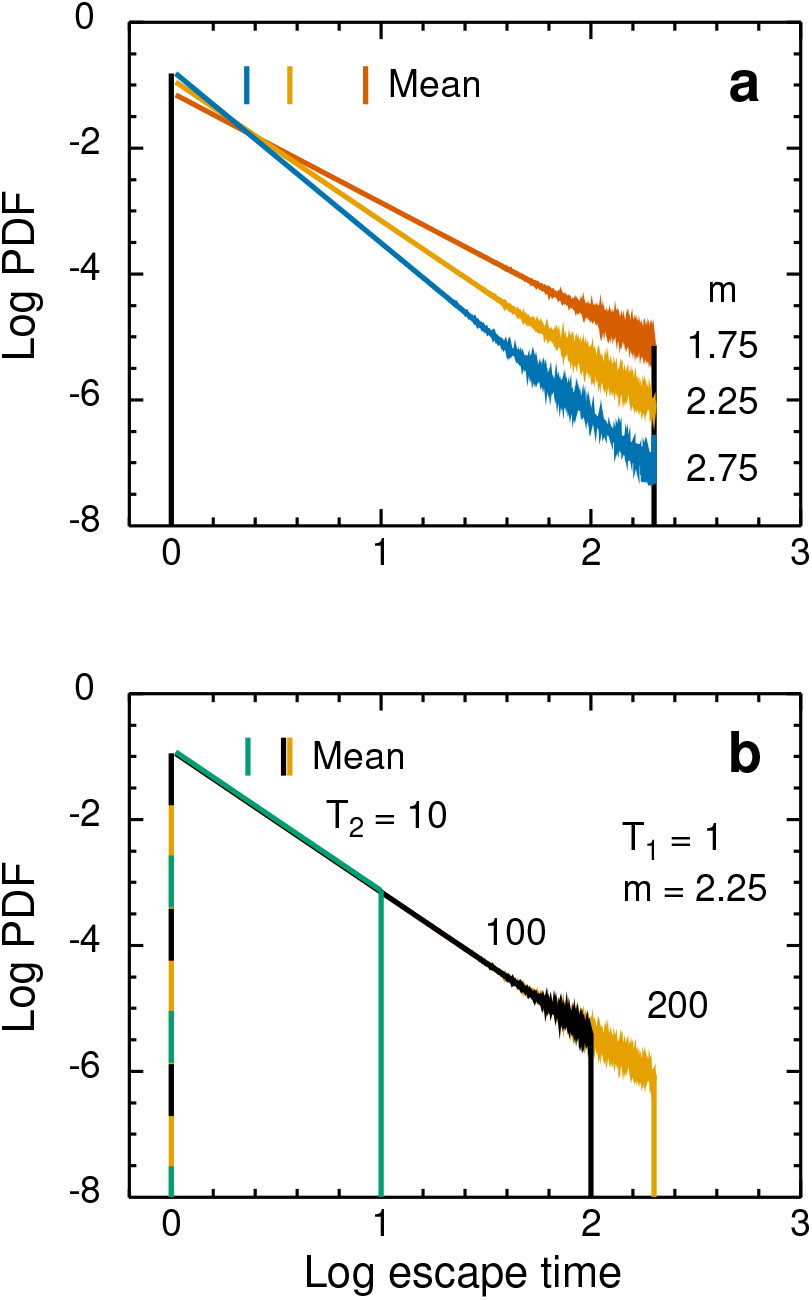
Log-log plots of truncated power law distributions for *(a)*, the specified values of the exponent *m* at fixed values of *T*_1_ = 1 and *T*_2_ = 200, and *(b)*, the specified values of *T*_2_ at constant *T*_1_ = 1 and *m* = 2.25. Vertical lines are means.

Further properties are given in Suppl. 2.2, and the case *m* = 1.

The experimental measurements are incomplete. The histogram for TetR (3) covers a range of escape times of roughly 0. 01 to 600 s, but the authors report that practical reasons lead to a 1 min delay between mixing and the start of the SPT measurements. The measurements on CRISPR-Cas9 interacting with a DNA curtain (6) would have a similar limitation if the measurement is of a tracer that is already in the DNA curtain, because one does not know how long it has already been there equilibrating. To avoid this problem, observations ought to be limited to tracers seen to enter the curtain region.

### From binding energies to observed escape times

We need to connect the distribution of observed escape times to the distribution of binding energies. This is done in two steps to better match the literature. First, we find the distribution of mean escape times corresponding to the distribution of binding energies. In the physics literature, this is the usual derivation of a power-law distribution of mean escape times. In the statistics literature, this is the derivation of the distribution of a function of a stochastic variable. Second, we find the distribution of observed escape times *τ* from the distribution of mean escape times *T*. In the physics literature, this is a very common procedure to express the escape times from a statistical distribution of traps in terms of a Laplace transform. In the statistics literature, this is a compound distribution, in which a parameter of the primary distribution is a stochastic quantity distributed according to the parameter distribution. (There is no consensus on nomenclature but “compound distribution” seems reasonable.) Both steps involve Boltzmann factors, a deterministic one in the first step and a stochastic one in the second.

We summarize some basic properties of the mean escape times *T*, which are related to Δ*E* by

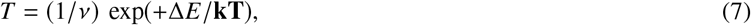

where *ν* is the frequency of escape attempts, Δ*E* is the absolute value of the free energy of binding, and boldface **kT** is the thermal energy. In the Fortran Monte Carlo calculations (Suppl. 1.1), *ν* is 1 per simulation time step. The escape probability per Monte Carlo time step is *Pesc =* 1/T; this probability generates the geometrical distribution of observed escape times in the Monte Carlo simulations. The Boltzmann factor for a tracer to be in the ith trap is

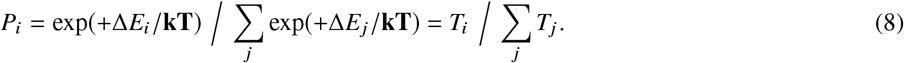

#### Step 1. From binding energies to mean escape times

What is the relation between the distribution of binding energies and the distribution of mean escape times, and what is the temperature dependence? The basic physical picture is that the system is at thermal equilibrium at the ambient temperature, except that the tracer is not necessarily equilibrated with the trap distribution. The distribution of binding energies is *ϕ*(Δ*E*, parameters), where Δ*E* has dimensions of energy and *ϕ* has dimensions of 1/energy, so that *ϕ*(Δ*E*)*d*Δ*E* is dimensionless. The parameters are constant values for the ambient temperature. The distribution of mean escape times depends on temperature only through the expression for the escape time *T* as a function of binding energy.

With this picture, the relation between the distributions is given by a standard transformation from statistics (22). The starting point is Eq. 7 with *ν* = 1. Then

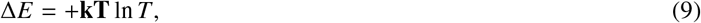

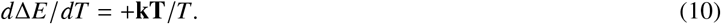

The distribution of mean escape times *g*(*T*) is

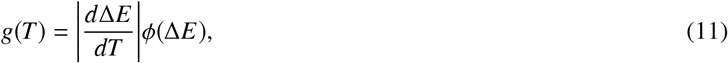

obtained by a mapping of CDFs in probability textbooks, or more formally as the Jacobian of the transformation. Then

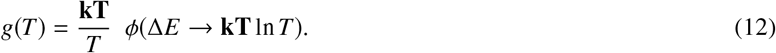

Here the mean escape time *T* is dimensionless, equal to the number of time steps *δt*, either Monte Carlo time steps (see subsection Conversion to Physical Units) or the duration of one frame in an SPT experiment.

What causes a power-law distribution of mean escape times? The simplest mechanism is Arrhenius escape from an exponential distribution of binding energies (3, 19, 23–25). Suppose that the binding energies are exponentially distributed on [0, ∞), so that the PDF is

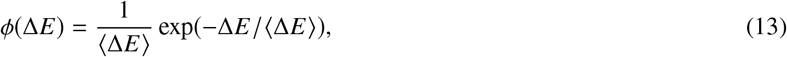

where 〈Δ*E*〉 is the mean binding energy. Then from Eq. (12)

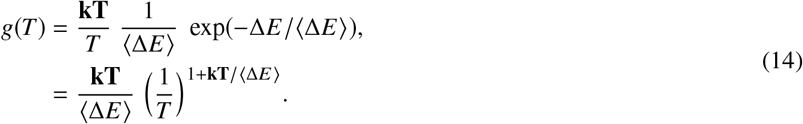

A pure exponential distribution of binding energies thus leads to a pure power-law distribution of mean escape times.

If we impose lower and upper bounds Δ*E*_1_ < Δ*E*_2_ on the binding energy, the derivation proceeds as before, except that the normalization integral of *ϕ*(Δ*E*) has finite limits so the distribution of mean escape times has limits *T*, = exp[+Δ*E_i_*,/**kT**], and the normalization constants are more complicated (Suppl. 3.1). The width parameter 〈Δ*E*〉 is replaced by a width parameter *θ*_trExp_ that is a complicated function of the upper and lower bounds. The distribution of *T*, Eq. S3–3, is then the product of the pure power law distribution Eq. 14, a truncation function Eq. S3–2, and the ratio of the width parameters. The resulting distribution is still power-law, as illustrated in Fig. 5 and derived in Suppl. 3.1.

**Figure 5:**
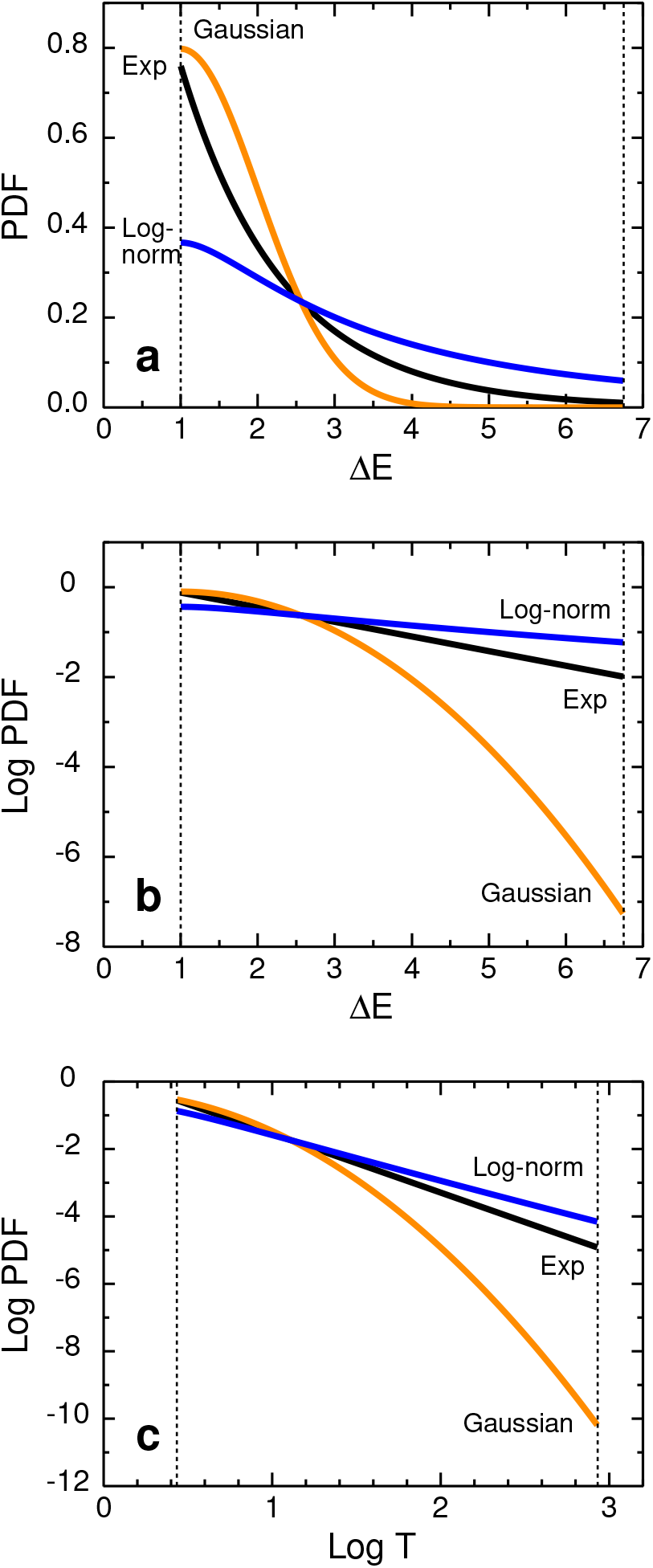
Truncated distributions of binding energies Δ*E* and the resulting distributions of mean escape times *T*. *(a)*, Linear PDFs of binding energies; *(b)*, logarithmic PDFs of binding energies; *(c)*, logarithmic PDFs of mean escape times. The PDFs in panels *b* and *c* are given as logarithms to show the tails clearly and to conform to practice in the genomics literature. Vertical lines show the truncation limits on the binding energy, Δ*E*_1_ = 1 and Δ*E*_2_ = 6.75, and the corresponding mean escape times *T*_1_ = 2.718 and *T*_2_ = 854. The limits were chosen to give an escape time distribution of width 2.5 decades, as in the experimental results from the Dahan lab. Analytical forms of the distributions are obtained in Suppl. 3.1. (1) Truncated exponential with θ_trExp_ = 4/3, **kT** = 1, so the power is 7/4. (2) Truncated Gaussian with *μ* = 1, *σ* = 1. (3) Truncated log-normal *μ* = 1, *σ* = 1. If the range included more of the peak of the log-normal distribution, there would be major nonlinearities in the corresponding part of the log-log plot of the PDF of the mean escape time.

A log-normal distribution of binding energies gives an approximately power-law distribution of escape times over a limited range, and the slope is tunable over a limited range via the temperature. A numerical example is shown in Fig. 5, the analytical form is given in Suppl. 3.1, and Fig. S3–1 *a* gives a numerical example of the effect of temperature.

Gaussian distributions of binding energies have a very limited ability to produce a power-law distribution of mean escape times. The Gaussian distribution is

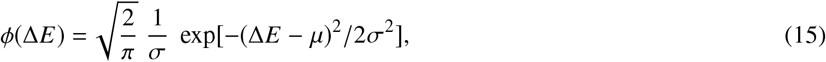

where *μ* is the mean and *σ* is the standard deviation. Proceeding as in the exponential case, we obtain a log-normal distribution of escape times

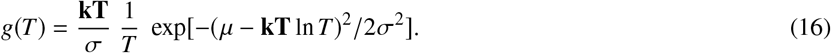

Here **kT**, *μ*, and *σ* have dimensions of energy, and the mean escape time *T* is dimensionless. In the numerical example of Fig. 5, *g*(*T*) from the Gaussian case is clearly far from a power law for the parameters chosen.

Log-normal distributions are known to have tails resembling power-law distributions over a limited range of the variable (26–28), but not under the conditions here. Take the logarithm of *g*(*T*) and rearrange:

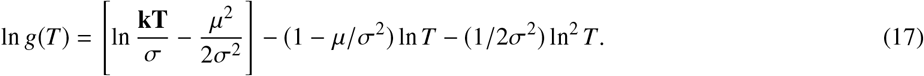

If we increase *σ* to decrease the quadratic term, we drive the coefficient of the linear term toward −1, which is fine if one is trying to reproduce Zipf’s law or 1/ *f* noise, but not useful here. Simultaneously matching the experimental slope and the range of linearity is difficult or impossible. Or, to put a more positive spin on it, the experimental signature of Arrhenius escape from Gaussian traps is a truncated power law with a slope near −1, potentially with a range of a few decades depending on the standard deviation of *f*(Δ*E*).

Gaussian distributions of binding energies are often used in the DNA literature (29, 30). These distributions are justified on the grounds that transcription factors typically bind around 10 nucleotides, range 5–31 (31), providing enough averaging to give an approximate Gaussian, whether the protein is reading hydrogen bonding or elastic interactions. This is roughly enough averaging. One algorithm to generate Gaussian random numbers is to generate 12 random numbers uniformly distributed between 0 and 1, sum them, and subtract 6, though this method gives such poor-quality results that it is highly unrecommended (16).

Some labs have used modified distributions such as an asymmetric Gaussian, for example, a saddle point expansion plus first-order correction (32), or Gaussian-like curves with tails that are defined numerically from experiment or bioinformatics (33), or defined analytically on the basis of an analogy between protein-protein binding and protein folding (34). But it is hard to see how these modifications could yield a TPL distribution over 2.5 decades in three different experimental systems.

Presumably a distinct mechanism is responsible for the observed TPL. One possibility is obstruction, say trapping in dead ends, but it seems difficult to make dead ends affecting diffusion on a scale < 25 nm out of DNA with a persistence length of 50 nm; “tightly bent” DNA structures would be required. Such structures were reviewed by Garcia et al. (35). Similarly, the fractal model of the nucleus (see for example Bancaud et al. (36)) is a coarse-grained model with beads of diameter 20-40 nm (37). Another possibility is the combined effects of multiple traps, recapture, and trap geometry, potentially important because a 25-nm segment of DNA includes 75 base pairs. Several groups have examined this effect (38–40) and Suppl. 7 of Ref. (3), but further work remains to be done. Other possibilities include viscoelasticity with crowding (41). and a nonequilibrium state produced by active cellular processes such as chromatin remodeling.

#### Step 2. From mean escape times to observed escape times

There are two distinct escape times in the problem, the observed escape time *τ* in an individual escape, and the mean escape time *T*. For escape from a binding site, *T* is determined by the binding energy, and for escape from a geometrical trap, by the obstacle geometry and dynamics. The distribution of *τ* is broader than the distribution of *T* due to the scatter in *τ* for a fixed value of *T*. The problem is, given a distribution of observed escape times *f*(*τ*), what is the distribution *g*(*T*) of mean escape times, assuming escape according to the Arrhenius law? Mathematical details are shown in Suppl. 3.2.

### Transient anomalous subdiffusion

In this section, we present results of Monte Carlo simulations of subdiffusion (Suppl. 1.1). Here *T* is taken to be the mean escape time, assumed to follow a TPL distribution as in Fig. 4. The actual escape time for each visit to a trap site is a random variate generated in the Monte Carlo program. In Fig. 6, panel *(a)* shows the effect of varying the exponent *m* at constant minimum and maximum times *T*_1_ = 1, *T*_2_ = 200. The curves are of similar shape but the value of log *t*(*cross*) changes with *m* as discussed in Suppl. 4.1. Panel *(b)* shows the effect of increasing *T*_1_ at constant *T*_2_ and *m*. As *T*_1_ is increased, the curves flatten and *D*(*t*) decreases. In the limit *T*_1_ = *T*_2_, the curve would be flat with *D*(*t*) = 1/*T*_2_. Panel *(c)* shows the effect of increasing *T*_2_ at constant *T*_1_ and *m*. As *T*_2_ increases, diffusion becomes more anomalous – the slope gets steeper – for a longer time. The top curve, *T*_2_ = *T*_1_ = 1, is the control; uniform traps yield normal diffusion at all times. (This curve is actually for *T*_1_ = 1, *T*_2_ = 1.01, not 1, so that the standard program could be used without modification.) Panel *(d)* shows that the trap concentration has a major effect on the anomalous exponent *α* at constant *T*_1_, *T*_2_, and *m*. As the fraction of traps increases, the curves grow steeper in the anomalous regime. The crossover time does not change much, so the change in slope is almost entirely due to the changes in *D*(0) and *D*(∞), Eqs. 19 and 23, with each nontrap site contributing *T* = 1 to the averages. Much, but not all, of the concentration dependence in the log *D*(*t*) curves can be removed by rescaling log *D*(*t*) as shown in Suppl. 4.2. In simulations, the limiting values of *D* are trivial to calculate from the TPL parameters. In experiments one ought to make good measurements of *D*(0) and *D*(∞), using fast and slow frame rates if necessary to get good values. The amount of averaging needed to show the TASD structure is discussed in Suppl. 4.3.

**Figure 6:**
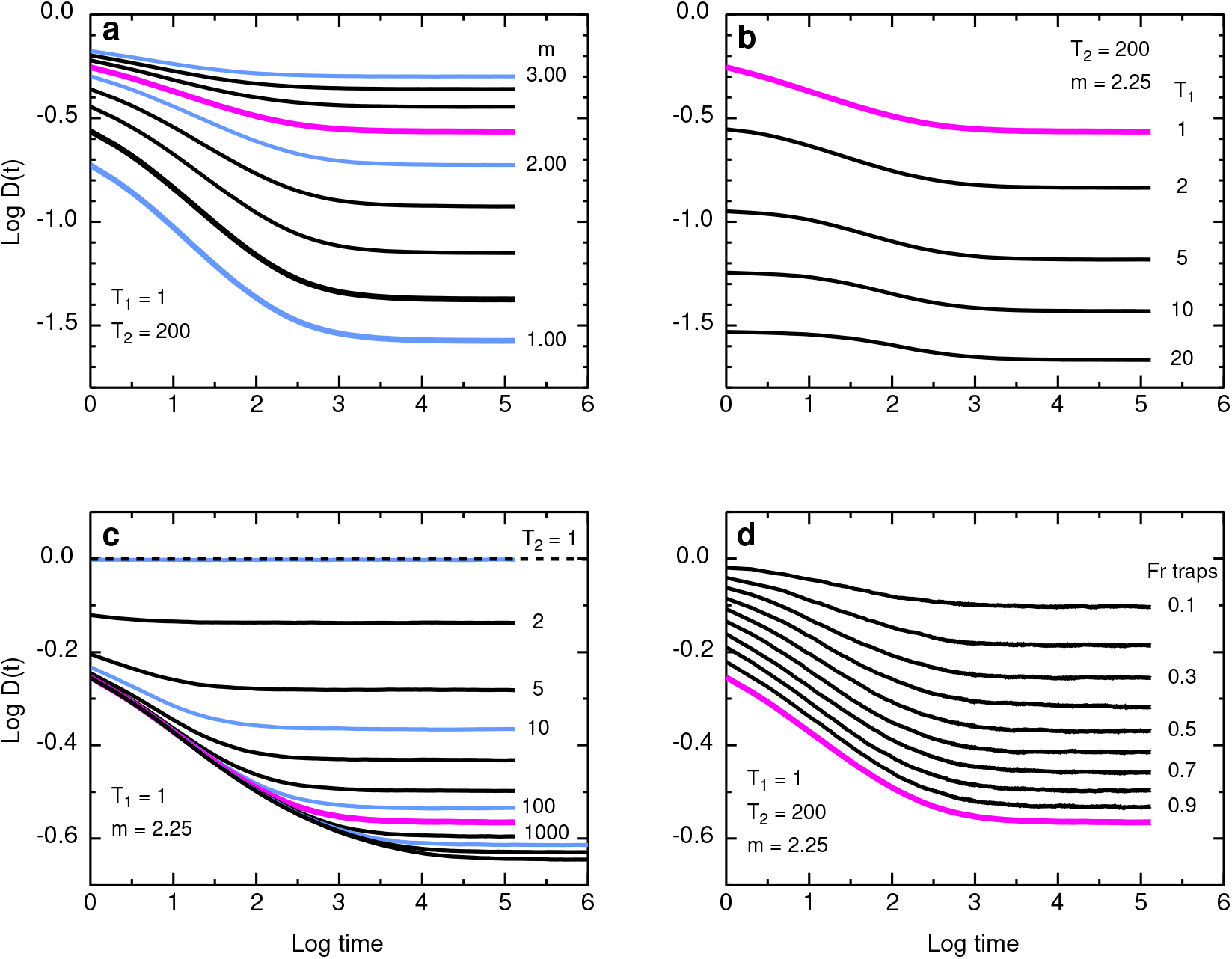
Log-log plots of *D*(*t*) versus *t*. Note the difference in the vertical scales for the first two panels and the last two. For comparison, the curve for the standard case *T*_1_ = 1, *T*_2_ = 200, *m* = 2.25, trap fraction 1, is shown in magenta in all panels. *(a)*, Vary *m* at constant *T*_1_ = 1, *T*_2_ = 200, trap fraction 1. Values of *m* are 1.00 to 3.00 in steps of 0.25. *(b)*, Vary *T*_1_ at constant *T*_2_ = 200, *m* = 2.25, fraction of traps 1, for the indicated values of *T*_1_. *(c)*, Vary *T*_2_ at constant *T*_1_ = 1, *m* = 2.25, trap fraction 1. Values of *T*_2_ are from 1 to 5000 in a 1, 2, 5 series. *(d)*, Vary fraction of traps at constant *T*_1_ = 1, *T*_2_ = 200, *m* = 2.25. Trap fractions are 0.1 to 1.0 in steps of 0.1.

Four parameters characterize *D*(*t*): *D*(0) and *D*(∞) describe the initial and final values, the slope *α* describes how anomalous the diffusion is, and t(cross) locates the transition from anomalous to normal diffusion. Here *D*(0) and *D*(∞) are obtained from simple formulas based on the TPL, but *α* and *t*(*cross*) are obtained from Monte Carlo calculations. Prediction of *α* and *t*(*cross*) will be discussed in future work.

The initial and final diffusion coefficients in a system with binding are given by a simple argument. The normalized diffusion coefficient is the product of the squared displacement for a move and the probability of that move. For a single step, the squared displacement is 1 in units of the lattice constant *ℓ*. For the first move, the probability is

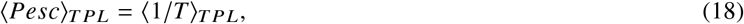

where the subscript indicates an average over the TPL distribution, so

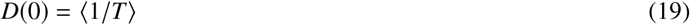

and from Eq. S2–7,

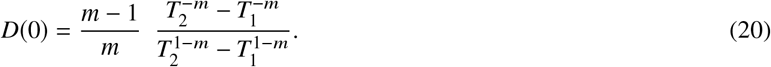

Here *D*(0) is normalized to 1 when all sites are nonbinding. For a move at long times, the probability is

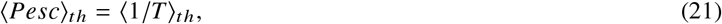

where the subscript *th* indicates a thermal average over the TPL distribution, so from Eq. S2–14,

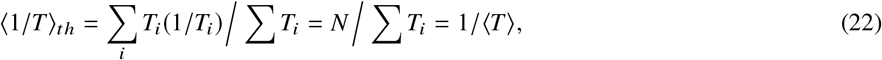

and

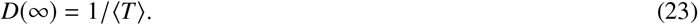

From Eq. S2–7,
>

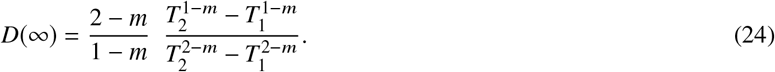

A more formal argument for Eq. 24 is given in the literature (42–44).

The results here for the 3D case are qualitatively similar to earlier results (1) for the 2D case. The main effect of dimensionality is through the probability that a diffusing particle revisits a site in a lattice model. For normal diffusion in an infinite system, the number of distinct sites visited (*DSV*) can be written as a function of number of steps *n*, as 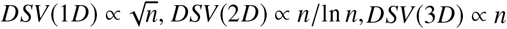. These are the leading terms; there are higher-order correction terms (45). For anomalous subdiffusion on an infinite fractal, the DSV depends on the so-called spectral or fracton dimension, which is not necessarily an integer. This dimension is one of the standard quantities used to characterize fractals (45).

### Conversion to physical units

A key question is how to translate Monte Carlo units into physical units, particularly how to translate *t*(*cross*) into clock time. The conversion is based on one of the fundamental equations of diffusion, in 3D

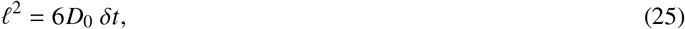

where *ℓ* is the lattice constant, *D*_0_ the diffusion coefficient in the absence of hindrances, and *δt* the unit of time. Choose any two and calculate the third.

In earlier work on pure obstructed diffusion in 2D membranes (46), the bilayer was taken to be on average a triangular lattice of lipids, *ℓ* was chosen to be the lipid diameter, and *D*_0_ was chosen as the value for the tracer in a pure lipid bilayer. The resulting *ℓ* is the characteristic time to diffuse the length of the bond joining adjacent lattice points.

The case of the nucleus is more complicated. Ultimately one would like to model all four hindrances to diffusion – binding, obstruction, viscoelasticity, and crowding – but at this stage it is appropriate to model binding in isolation, and take the other hindrances into account approximately through *D*_0_. We choose *D*_0_ to be the value from FRAP experiments on GFP in the nucleus of H1299 human large cell lung carcinoma cells (47). This is a long-range diffusion coefficent for a protein that is presumably nonbinding. The value is *D*_0_ = 41.6 *μ*m^2^/s, here rounded off to 40 *μ*m^2^/s. What value to use for *ℓ*? Analogy with the membrane case suggests choosing *ℓ* to be the size of a transcription factor binding site, roughly 10 base pairs (31), so 3 nm. Then *δt* = *ℓ*^2^/6*D*_0_ = 37.5 ns, which is very poorly matched to the time resolution of the SPT experiments. A single SPT time step of 5 ms would require over 128K Monte Carlo steps. A better choice is to take *ℓ* to be the SPT spatial resolution of 25 nm, so that *δt* = 2.6 *μ*s. This is still not well-matched to the SPT experiment; a single SPT time step corresponds to 1920 Monte Carlo time steps, and a escape time of 5 s requires 1.8M Monte Carlo time steps. But any further increase in *ℓ* would make the Monte Carlo resolution less than the experimental resolution. Better to spend computer time on longer simulations than to throw away hard-earned experimental spatial resolution. With this choice of *ℓ*, the Monte Carlo calculations are done on a simple cubic lattice of resels, resolution elements, which seems appropriate.

### Nonequilibrium requirement

A fundamental property of a model based on time-independent (quenched) energy traps is that it is highly sensitive to the initial conditions, specifically the time since the interaction of the tracer with the traps is turned on. In the nonequilibrium case, fresh tracer diffuses with a time-dependent diffusion coefficient *D*(*t*) as shown in Fig. 1 *d*. In the equilibrium case, stale tracer diffuses with a small constant diffusion coefficient *D*(∞), determined largely by the escape time from the deepest traps.

In the physics literature, this property is called “aging,” the dependence of diffusion on the time elapsed since the initial preparation of the system (24). In the case of DNA binding, this property is another way of stating the idea that the initial period of anomalous subdiffusion represents the search of the diffusing species for the deepest traps, that is, for its biological target.

The experimental work gives a variety of situations. In the nucleus, the interaction is turned on by entry of the tracer into the nucleus, or by biochemical activation. For CRISPR-Cas9 binding to a DNA curtain (6), the interaction is turned on by 3D diffusion from bulk solution to the DNA curtain, and the clock is stopped if the tracer leaves the curtain and re-enters bulk solution.

Normanno et al. (3) reported a time of 1 min between mixing and the start of observations, a significant delay. Modeling of delay effects is discussed in Suppl. 4.4, but the main implication of the modeling is that experimental data is needed. Photoactivation of the DNA-binding species would be very useful experimentally, to reduce the delay and to define “*t* = 0” precisely for ensemble averaging of measured trajectories. One approach to photoactivation is caging (48, 49), but the caging must be of the binding domain, not the functional domain. The “CRISPR-plus” of Jain et al. (50) meets this requirement. Another approach is (a subset of) optogenetics, (51–53). For the application here, a unimolecular reaction is better than a bimolecular one, that is, uncaging or conformational change is preferable to dimerization. A unimolecular reaction reduces the delay time between the light pulse and activation, and reduces the scatter in delay time. The magnitude of the delay time is critical, and we require much shorter delay times than in many applications. Optogenetics workers often measure the time course by protein synthesis, well downstream of the binding events considered here, with a time scale of hours rather than seconds, so measurements with higher time resolution are needed. Furthermore, we need fast switching on, but slow switching off, if any. Despite all these caveats, optogenetics work on a transcription factor showed a useful increase in speed even though the reaction is bimolecular. Motta-Mena et al. (54) developed an inducible promoter based on the EL222 light-activated promoter. The promoter is smallish, 33 kDa, and modeling of their protein expression experiments gave a delay time of 5 s or less; this delay time includes the entire process from search to protein expression. Other work characterized an EL222 mutant with a stable photoactivated state. It is encouraging that a protein designed and optimized for other types of experiments has properties that would be useful in work on binding kinetics. Light-regulated gene transcription was reviewed recently (55).

We can illustrate this effect quantitatively by Monte Carlo calculations in which the tracer takes a prescribed number of equilibration steps before log *D*(*t*) is recorded. (See (2) for the 2D case.) For an equilibration time of 0, we recover the usual TASD results. For the largest equilibration time, diffusion is normal at all times but slow, with *D* = *D*(∞). Intermediate equilibration times flatten the initial part of the *D*(*t*) curve, as shown in Fig. 7. The theoretical importance of this result is that one can go smoothly from pure TASD to slow pure normal diffusion by tuning a single well-defined parameter. The experimental importance is that the changes in shape at small times are the signature of partial equilibration of the tracer with the binding sites, for example, during the delay between mixing and the start of SPT observations. Importantly, no such changes occur for pure obstructed diffusion. The plot of log *D* versus log *t* is independent of initial delay time (1).

**Figure 7:**
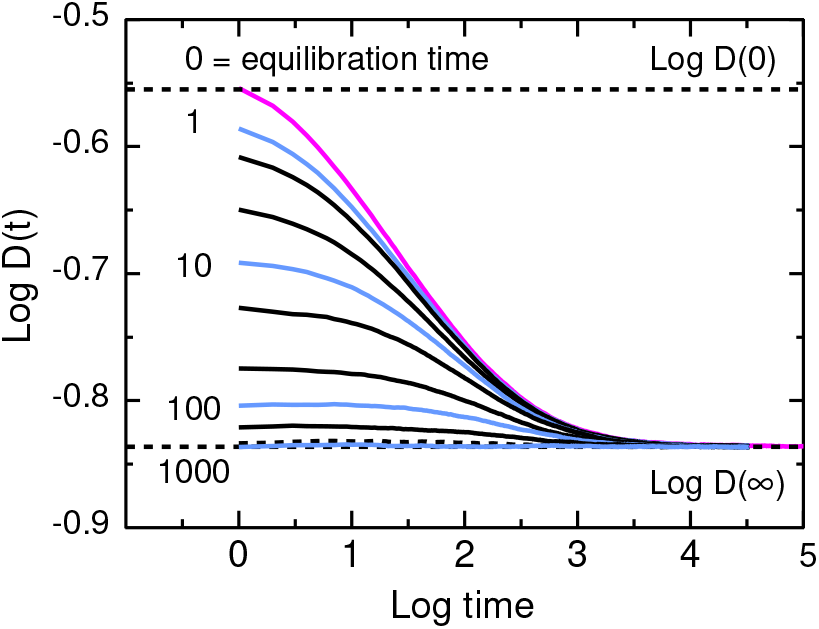
Annealing. Change in shape of log *D*(*t*) versus log *t* for various equilibration times: 0, 1, 2, 5, 10, 20, 50, 100, 200, 500, 1000, with *T*_1_ = 2, *T*_2_ = 200, *m* = 2.25, trap fraction 1. In general, significant equilibration effects occur for equilibration times < *t*(*cross*).

To specify the nonequilibrium state more precisely: The tracer is in local thermodynamic equilibrium, so its probability of being in a trap is given by a Boltzmann factor, as in Eq. 8. But the tracer is not in global equilibrium with the population of traps, and TASD reveals this global equilibration process, a process that requires the tracer to sample the population of traps adequately, as will be discussed in future work.

## DISCUSSION

How does the model presented here connect to results in the physics and biophysics literature?

### Connections to models of anomalous subdiffusion in the physics literature

Several different physical mechanisms can lead to anomalous subdiffusion, pure or transient. See, for example, (56) and recent reviews such as (24, 57–60).

a. *Nonequilibrium trapping in energy traps*. As already discussed, the tracer must not be in equilibrium with the system of traps. If it is equilibrated, it diffuses normally, at a rate dominated by the escape time from the deeper traps.
b. *Obstruction*. Obstruction can lead to pure or transient anomalous subdiffusion (17). There is no dependence on equilibration time but obstructed diffusion is highly sensitive to tracer size, as discussed in the next subsection.
c. *Anticorrelated motion*. In the physics literature, this category has often been taken to be fractional Brownian motion as defined by Mandelbrot and Van Ness (61) in terms of a correlation function of extremely macroscopic inspiration, Hurst’s classic studies of flooding on the Nile. A more microscopic starting point for cell biophysics is viscoelasticity (58, 62, 63).
d. *All of the above*. These processes are nonexclusive and can operate in parallel. In a cell, it would be much more surprising to have one of the processes dominant than to have all three act simultaneously. The obvious example is that chromatin can act as an obstacle and as a set of binding sites. For examples of how to sort out anomalous subdiffusion mechanisms, see Golan and Sherman (64) for the T-cell plasma membrane; Kepten et al. (65) for diffusion of telomeres; Szymanski and Weiss (66) for a model crowded system; and Thiel et al. (67) for modeling.

A series of experiments and simulations by Izeddin et al. (68) addressed the question of physics models by experiment and simulations. The experiments used PALM SPT to examine motion of two nuclear factors in U2OS human osteosarcoma cells. Trajectories were analyzed by plots of mean-square displacement versus time, histograms of step sizes, and histograms of the angles between consecutive displacements at various time lags as a test for anticorrelation on various length scales. Four proteins were used, all with the photoconvertible label Dendra2: (1) Free Dendra2. It is structurally similar to GFP, and is assumed to be freely diffusing because GFP has no known binding partners in mammalian cells. (2) Dendra2 fused to the proto-oncogene c-Myc, which binds directly to DNA. Fast, slow, and immobile populations were observed, and the angular distribution of moves was approximately uniform. (3) Dendra2 fused to the cyclin T1 subunit of P-TEFb, which binds to the transcription machinery. Motion was subdiffusive with α ~ 0.6, and the angular distribution of moves favored reversal. (4) Dendra2 fused to histone H2B was bound, and the angular distribution favored reversal. Results were compared to a variety of models based on the physics literature, and interpreted in terms of target-searching strategies. For c-Myc, the search involved global exploration of space, which leaves sites unvisited. For P-TEPb, the search involved local (compact) exploration of space, and thus a redundant search.

### Connection to models of the nucleus in the biophysics literature

Several interpretations are possible of the diffusion model presented here.

a. *No interpretation is needed*. A truncated power-law distribution of mean escape times necessarily implies transient anomalous subdiffusion, independent of the microscopic mechanism producing the escape times. But it is useful to go beyond this to ask, what happens during the escape time?
b. *Binding*. The diffusing particle is bound to an immobile site on the DNA, escapes, and then diffuses freely or at the rate set by pure obstruction. This model is the one discussed in the most detail here.
c. *Facilitated diffusion*. To reduce a vast literature to a single paragraph, the key problem in DNA binding is searching. How does a DNA-binding protein find its biological target fast enough among the huge number of potential binding sites? The problem is solved by the classic Berg-von Hippel facilitated diffusion model (7, 8), in which the search is speeded by switching between 3D and 1D diffusion. A major issue within the facilitated diffusion model is the speed-stability paradox, involving the energy landscape of the protein-DNA interaction. A smooth landscape is required for rapid motion, but a rugged landscape is required for specific binding. A solution to this problem is the two-state protein model, in which the protein can exist in recognition and search states (29). Recent reviews include the very succinct Ref. (69), more detailed discussions in Refs. (30, 70, 71), and a physics perspective in Ref. (72), but see Ref. (73) for an argument that the speed-selectivity paradox is not real. The energy landscape can lead to 1D subdiffusion by the same mechanism as discussed here: TASD until the potential averages out, as discussed by Barbi et al. (74, 75). The simplest mapping between facilitated diffusion and the model presented here identifies the escape time from a trap as a period of 1D diffusion, and diffusion between traps as a period of 3D diffusion, as suggested by Normanno et al. (3). This mapping is consistent with the resolution of the experiments. Estimates of the 1D diffusion length vary among systems; Barbi et al. (74) used a value of 170 base pairs. The SPT spatial resolution is 25 nm or 75 base pairs (1 base pair = 0.332 nm), so a period of 1D diffusion of 170 base pairs is below the 1D detection limit of 3 points for a straight line. The persistence length of DNA is 50 nm (15) so the DNA in one resolution element could be slightly curved, but not coiled unless bound in a nucleosome. A related model comes from recent work in the Xiao lab on SPT of RNA polymerase (RNAP) in *E. coli* cells [Bettridge et al. (76, 77), Bohrer et al. (78)]. Labeled RNAP was photoactivated one molecule at a time, its motion was tracked, and the trajectories were analyzed by hidden Markov modeling using the algorithm of Persson et al. (79). In preliminary results, three distinct diffusive modes were found, and were interpreted as DNA-bound RNAP, RNAP rapidly associating and dissociating with DNA, and freely diffusing RNAP. The authors argued that the system is not in a steady state; the occupation of the states is not that predicted from the escape times. They suggested that nonspecific binding is the rate-limiting step in promoter search. Their model maps directly to a version of my model extended to include nonbinding sites. The experimental distribution of escape times will be of interest. Their DNA-bound state corresponds to binding to the target site or possibly the deepest false-positive sites; this ambiguity leads to the question, what is the gap (in binding energies or in escape times) between the target site and the deepest false-positive sites? Sheinman et al. (30) discussed this question in detail, obtaining the energy of the deepest false-positive site from extreme value statistics, and distinguishing the gapped, marginally gapped, and non-gapped cases. Another possibility could be called nested TASD. Here the first level is 1D TASD due to the detailed DNA sequence, resulting from variations in either the H bond binding energy (74, 75) or the elastic energy of DNA bent by protein (10). This diffusion crosses over to normal when the DNA-binding protein has sampled the DNA sufficiently. The second level is TASD due to the distribution of escape times in the 1D diffusion state. The resolution of the experiment determines how much of this can be observed. Nested TASD can be considered a multilevel equilibration process.
d. *Obstruction and geometrical traps*. Diffusion in the nucleus is likely to be affected by obstruction, especially by geometrical trapping in dead ends (see Suppl. 5). Obviously the geometrical traps must be large enough for the tracer to enter, as in size exclusion chromatography. If the trap size is below the resolution of the SPT measurement, the tracer will appear to be stationary. If motion in the trap can be resolved, the well-known form of 〈*r*^2^(*t*)〉 for confined motion (80) will be observed, with eventual escape to free diffusion. This form is distinct from the form for anomalous subdiffusion, as shown in Fig. 8. It is informative to make two plots, 〈*r*^2^〉 versus *t*, and log *D* versus log *t*. In their work on TetR diffusion in the nucleus, Normanno et al. (3) observed some trajectories consistent with pure confinement, with time scales < 1 ms and confinement radii in the range of 0.5–1 *μ*m. (See their Supplement Fig. 2, Confined diffusion analysis, and their Supplement Note 4, Single-particle tracking analysis.)

**Figure 8:**
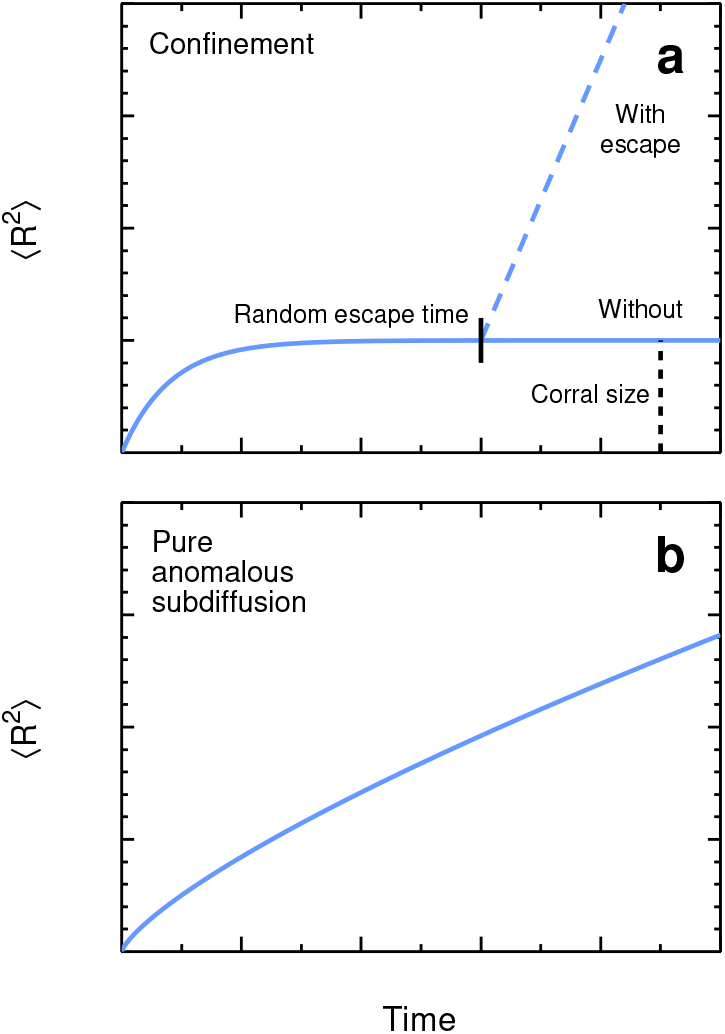
Cartoon of 〈*r*^2^〉 versus time for geometrical trapping and for pure anomalous subdiffusion. *(a)*. Pure confinement, and transient confinement with eventual escape and free diffusion. The plateau is a measure of the corral size, and the escape time is random. *(b)*. Pure anomalous subdiffusion. There is no plateau.

For rigid immobile obstacles, there is a well-defined percolation threshold. The experimental signature of this case is that diffusion is highly sensitive to the size of the tracer. The factor controlling diffusion is not the area fraction of obstacles but the excluded area fraction, which is highly sensitive to tracer size [for 2D immobile obstacles, (81)]. Benichou et al. (11) modeled chromatin as a 3D percolating cluster at the threshold. They argued that on a large scale, chromatin is effectively branched as a result of intersegmental transfer of the diffusing species.

For soft or mobile obstacles, there is no permanent obstruction. The escape times depend in part on obstacle dynamics, such as mobility, dissociation rates, or rates of conformational fluctuations. Chow and Skolnick (82) present a coarse-grained model of gated diffusion of LacI in the *E. coli* nucleoid, in which LacI moves from cage to cage when the DNA wall moves. Diffusion may still be highly sensitive to size as a result of the lower probability of large fluctuations. High concentrations of mobile obstacles lead to a glassy state. For a model of the nucleus based on glassy dynamics, see Kang et al. (83).

Experiments using various nonbinding probes would be useful. Ideally one would use “scalable tracers,” in which the size of the tracer is varied but with constant surface properties, as discussed in Suppl. 5 and in detail in (84). One of the more speculative of these – a research project in itself – is “Sloppy Quantum Dots,” quantum dots synthesized with loose control over the dot radius but a constant thickness of the surrounding inert shell, to give a preparation with a broad range of sizes, inherently color-coded for size by the fluorescence. Less adventurously, a mixture of commercial quantum dots of different sizes could be used. The Lidke group has reported SPT of multicolor quantum dot mixtures (85). Genetically coded scalable tracers based on intrinsically fluorescent proteins would be highly advantageous. GFP oligomers meet the requirements of constant surface properties and interactions, but not the requirement for constant shape, because they are cylinders of constant diameter but different lengths. Several labs have reported experiments on these oligomers. Bancaud et al. (36) analyzed diffusion in terms of their fractal model. Dross et al. (86) measured diffusion of eGFP oligomers in four cell lines by careful FCS. Diffusion in bulk solution was consistent with a rod-like structure, and diffusion in the nucleus was independent of chromatin density. The Rippe lab (87, 88) measured diffusion of various GFP oligomers in the nucleus using multiscale fluorescence cross-correlation spectroscopy. Very promising recent work from the Holt lab has developed genetically encoded spherical tracers, called GEMs (Genetically Encoded Multimeric nanoparticles) (89–91), that meet the requirements for scalability. These tracers are based on natural proteins that self-assemble into spherical particles. The GEM DNA codes for the self-assembling monomer plus an intrinsically fluorescent protein plus a cellular localization signal if needed. Each monomer has one intrinsically fluorescent group attached, so the fluorescence is bright and the surface is for the most part constant. Tracer diameters are 16, 37, 100, and 200 nm at present. The self-assembling proteins used include encapsulins, which are proteins of bacteria and archaea that naturally encapsulate protein cargo (92). This approach may be extended using computational design of self-assembling cagelike proteins (93).

## CONCLUSIONS

The model has several implications for experiments and the interpretation of experiments.

a. *Search*. On the scale of the lin-lin plots, the anomalous regime seems small, but its existence and properties are clearly revealed in log-log plots of *D*(*t*) as shown in Fig. 1, so this analysis is worth doing as an examination of the search process.
b. *Escape time distributions*. We have shown the relation between the observed escape times and the mean escape times for a TPL distribution of mean escape times. The distribution of observed escape times is broadened by the statistics of escape. The mean escape time distribution is static; the observed distribution is static plus dynamic. For binding, the mean escape time distribution will eventually be predictable from genomics, though this distribution of binding energies must be corrected for the physical accessibility of sites to a tracer of given size and shape. For obstruction, constructing the escape time distribution requires a geometric model of chromatin.
c. *Limiting measurements*. The values of *D*(0) and *D*(∞) capture much of the dependence of *D*(*t*) on the TPL parameters. These values are trivial to calculate from the parameters, and essential to measure experimentally, using fast and slow frame rates if necessary.
d. *Experimental controls*. As claimed in the subsection on models of anomalous subdiffusion from the physics literature, binding, obstruction, viscoelasticity, and crowding may all affect diffusion in a cell simultaneously. Sorting out these mechanisms out requires a variety of experimental controls even more extensive than usual. As a baseline, *D* in buffer ought to be measured for each probe. For comparisons among cell lines and laboratories, *D* of eGFP ought to be measured. Two types of controls are needed for nonspecific binding by the DNA-binding species of interest: “specific nonspecific” binding, in which the recognition site binds to DNA sequences that only partially match the true target site; and “nonspecific nonspecific” binding, in which nonrecognition sites bind to DNA, say through electrostatic interactions. Natural selection may have reduced nonspecific binding, as discussed by Qian and Kussell (94), and Buchanan (95). The probe for “specific nonspecific” binding ought to be the DNA-binding species of interest with the binding site inactivated as unobtrusively as possible. For example, the pioneering single-particle measurements of Elf et al. (96) on transcription factor dynamics used LacI (360 amino acids) with the DNA-binding domain deleted (41 amino acids); the fusion to Venus fluorescent protein (238 amino acids) was most likely a much larger perturbation. Elf and Barkefors (97) reviewed single-molecule kinetic measurements in live cells, with an extensive discussion of labels and perturbations from labeling. The comparison of nonbinding and binding tracers is essential to be able to distinguish TASD due to obstruction from TASD due to binding. A general problem in diffusion measurements in complex systems is that a measurement of *D* or *D*(*t*) is not uniquely invertible. By an appropriate choice of parameters, the three anomalous subdiffusion mechanisms mentioned above can presumably be tuned to produce very similar curves, and the mechanisms may well operate in parallel. Despite this limitation, diffusion measurements are important to biophysicists because diffusion is important to cells; diffusion is a means of transport that uses thermal energy, not metabolic energy – as Pederson (98) put it, running off the energy of the Big Bang. Histograms of escape times for nonbinding tracers would help to characterize confinement events due to obstruction and apparent confinement events due to fluctuations in pure random walks, in the context of the experimental conditions and the criteria used to define confinement. These criteria ought to be described in detail, as in Gorman et al. (99) for example. Nonspecific binding can be characterized by comparing the escape time histograms for the DNA-binding tracer and the corresponding nonbinding control. The controls for obstruction ought to be done experimentally. Modeling obstruction requires a model of chromatin organization in the nucleus, a complicated and unsettled issue at best, requiring multiscale modeling. See for example (12–15, 100).
e. *Sub-resel dynamics*. In the experiments from the Dahan lab, the resolution element was significantly larger than a transciption factor binding site, so it is necessary to consider the dynamics of the DNA-binding species within a resel. Experimental data would be a useful constraint, particularly a histogram of the escape time from a single resel for a sample of distinct resels. These histograms ought to be reported for the DNA-binding species, the nonbinding analogs, and eGFP or another general calibration protein.
f. *Effect of deep traps*. Eliminating deep traps may affect diffusion and kinetics. First, false-positive sites affect diffusion, and the effect is greater the more similar they are to the target sequence. As a result, in bioinformatics searches for binding sites, the occurrence and frequency of false-positive sites are important results, not annoying artifacts. (For a general discussion of weak binding sites on DNA, see Tanay (101).) The search process in vivo might be speeded by (re)designing the genome to eliminate false-positive sites, that is, by separating the target sequences in affinity space. A mild form of this effect was found in prokaryotic genomes (102). Second, overexpression of the diffusing species affects diffusion. The level of expression is potentially an uncontrolled variable affecting TASD, or a means of modulating and studying TASD. In their work on TetR in U2OS cells, Normanno et al. (3) did a key experiment to address (and refute for their system) this point. They tracked labeled TetR with and without 1000-fold excess of unlabeled TetR. They saw no effect. The traps were not saturable, at least at this level of excess. Perhaps the traps were highly abundant geometric traps. This sort of test ought to be standard practice. As these authors point out, quantifying the number of binding sites in the nucleus would be very useful. This effect of eliminating deep binding sites can be approximated very simply if the concentrations of binding sites are known. If *N* unlabeled tracers are added, assume that they fill the *N* deepest sites, in effect reducing the upper limit *T*_2_ on the mean escape time and making diffusion less anomalous for a shorter time as shown in Fig. 6. This description is somewhat fictitious in that it assumes that *N* unlabeled tracers at zero temperature fill the deepest traps permanently, and then one labeled tracer at ambient temperature diffuses, entering into and escaping from the remaining traps. But the picture captures enough of the physics simply enough to be a named model, the step approximation. A more complete treatment includes the chemical potential to allow thermal excitation of all bound tracers. Such a treatment is a standard result in the field of diffusion in inhomogeneous catalyst pellets (103–105). Models of DNA binding that include the chemical potential are called “biophysical” or “thermodynamic” or “statistical thermodynamic.” See for example (101, 106–108). A rigorous approach takes into account the binding equilibria of multiple species to multiple sites, as in the program of Wasson and Hartemink (109).
g. *Applications of model*. The TASD model provides a simple Monte Carlo prediction of *D*(*t*) based on SPT measurements, suitable for use in larger-scale modeling of FRAP, FCS, and kinetics. It would be interesting to do FRAP or FCS on the same system as this SPT work, with the corresponding Monte Carlo simulations.

As expected, the Monte Carlo simulations presented here show that a truncated power-law distribution of escape times leads to transient anomalous subdiffusion. We have described the parameters characterizing each, and begun to examine the connection between the two sets of parameters. We have discussed the biological interpretation, and the experiments and controls suggested by modeling. Further work will examine the detailed connection of the TPL and TASD parameters, and the basis for a TPL distribution of escape times.

## ACKNOWLEDGMENTS

The early parts of this work were supported by NIH grant GM-038133.

## SUPPLEMENTARY MATERIAL

An online supplement to this article can be found by visiting BJ Online at http://www.biophysj.org.

### 1 METHODS

#### 1.1 Fortran Monte Carlo calculations

Here we give details of the Fortran Monte Carlo calculations and discuss approaches to such simulations. In the calculations, tracers carry out a random walk among trap sites, and the mean-square displacement is found as a function of time.

The calculations were carried out on a simple cubic lattice with periodic boundary conditions, with the standard system size of 128^3^ chosen so that the periodic boundary conditions do not affect the time dependence of the number of distinct sites visited. (This choice will be important in future work on the problem.) In each run, calculations were carried out for 500 grids, and 2000 tracers per grid. In each grid, a random TPL mean escape time was generated for each lattice site. (A more efficient approach would generate mean escape times on-the-fly, only when a site was actually visited, and retain the mean escape time at that site for all walks on that grid.) Every site is occupied by a trap unless specified otherwise. Tracers were started at random positions. This choice yields a highly nonequilibrium initial state; thermal equilibrium initial positions would lead to much different behavior, as discussed in the text. A minimum of 128K time steps were used, and sometimes as many as 2M (1K = 1024, 1M = 1024^2^). Data was collected in a block logarithmic pattern: from 1 to 1K in steps of 1; from 1K to 4K in steps of 4, from 4K to 16K in steps of 16, and so forth (110).

The simplest algorithm was used for moves: At each time step, one attempt to leave the current site is made. (Generate a random variate *r* uniformly distributed on (0,1). Move if *r* < *P_esc_* = 1/*T*, where *P_esc_* is the escape probability and *T* is the mean escape time.) The model assumes each trap is isolated, so the probability of escape depends only on the depth of the current trap, not on the depth of the destination site. (In other words, the motion is from trap to ground state to trap, not directly from trap to trap.) One can therefore test for escape at each time step, and generate the random destination only when escape occurs. Note that each move requires at least one time step, so that in the limit of traps of zero depth, 〈*r*^2^〉 = *t* and *D*(*t*) = 1 for all *t*.

An approach in the spirit of the “N-fold way” of the physics literature (111) ought to be more efficient. Here, instead of generating a whole random sequence of failures before a move, the program generates a random escape time from a geometric distribution with the required mean, increments the clock by that time, and moves the tracer to a random destination. But it is necessary to fill in all the observed quantities at all the intermediate time steps, and the intermediate time steps involve the block logarithmic sampling times. The run time was not reduced enough to justify the increased complexity.

The ran2 random number generator of Press et al. (112) was used, slightly modified. To avoid unnecessary overhead from branching and function calls at every time step, the initialization branch was separated and done in the program initialization, and the branch to generate the random numbers was included explicitly wherever it was used.

#### 1.2 Fitting procedure for Fortran Monte Carlo data

Ultimately the analysis of the Monte Carlo data ought to be done by fitting to the analytical form of *D*(*t*), but the form is not yet known. Instead, we plot log *D*(*t*) = log〈*r*^2^〉/*t* as a function of log *t*, and find the midpoint log *t*(1/2) from the analytic values of log *D*(0) and log *D*(∞). We then do a linear fit about that point, starting with 3 to 5 points on each side of the midpoint, and then selecting the linear region interactively by eye. The slope of the best-fit line yields the exponent, and the intersection gives the crossover time. In my experience, the method is far more objective than one might guess, the comments of Ellery et al. (113) notwithstanding.

#### 1.3 Mathematica Monte Carlo calculations

It was useful to run some Monte Carlo simulations in Mathematica to relate the observed escape times *τ* to the mean escape times *T* for a given distribution. These simulations were run alongside the derivations and the Fortran Monte Carlo simulations, and were a useful consistency check, able to find errors in numerical factors. (Similarly, I’d suggest that SPT experimentalists run Monte Carlo random walk simulations in parallel to their experiments. Anything done to the experimental data also ought to be done to simulated data.) These calculations were done for the uniform distribution as the simplest test, and the TPL for the application.

The method of histogram binning is an important practical matter in these calculations, especially for a power-law distribution of mean escape times. The histograms are much better-looking and easier to test if binning is done on what is called a “power scale” or a “log scale” (19). That is, for a typical run here, *root* = 2500^1/200^, *bins* = *root^i^*, for *i* = −200,200 to give a range of 1/2500 to 2500, with bins of equal width in a log plot.

The default Mathematica V11.2.0 pseudorandom number generator was used (the ExtendedCA generator = extended cellular automaton generator). This generator is discussed in online documentation (114) but I have not found an account of the method and its testing in the peer-reviewed literature. Apparently extensive testing was carried out (115, p. 231). There is a useful discussion online of the quality of the generator (116).

##### 1.3.1 Simulations for the uniform distribution

We begin with the simplest nontrivial example, a uniform distribution of the mean escape time *T* on the interval [*T*_1_, *T*_2_].

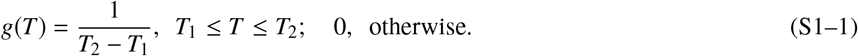

We insert this into S3-17 to obtain

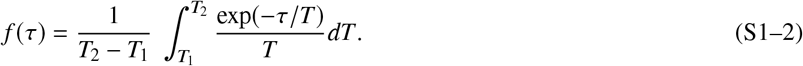

so that

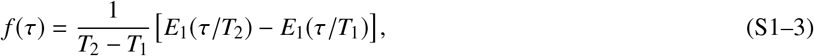

where *E*_1_(*x*) is the exponential integral.

Monte Carlo runs were carried out in Mathematica. Simply generate a set of mean escape times *T*, here 10^4^ uniform random variates on the interval [1,125], and for each *T*, generate 10^4^ exponential random variates *τ* with mean equal to *T*. Compile power-scale histograms for *T* and *τ*. Results are shown in Fig. S1–1. The Monte Carlo results agree well with the analytical results on the scale of a log-log plot of the entire data set. A more stringent test is plotting the ratio of the Monte Carlo values to the analytical values. Deviations are obvious, but these are low-frequency noise, not systematic error, as shown by the variation in the noise among three independent simulations. (Data not shown but the corresponding plots are shown for the TPL case.)

**Figure S1–1:**
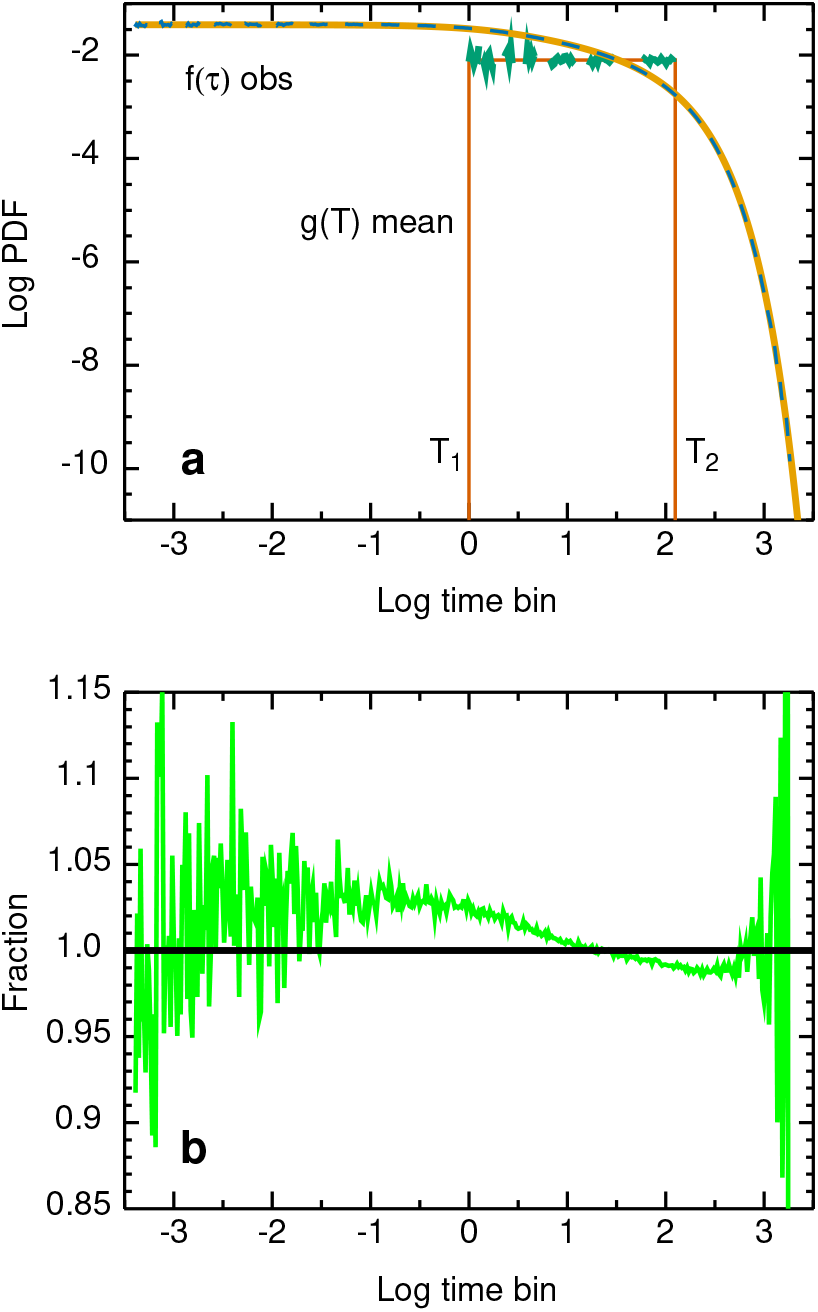
Results of Monte Carlo simulations in Mathematica for the uniform distribution on [1, 125]. *(a)*, Histograms of the TPL distribution of mean escape times *g*(*T*), analytical (red) and Monte Carlo (dashed blue-green), and the resulting distribution of observed times *f*(*τ*), analytical (orange) and Monte Carlo (dashed blue). The uniform histogram of *T* is noisy at small values of log *T* because the bins are smallest there. *(b)*, Ratio of Monte Carlo results to analytical results at the bin center (taken to be the geometric mean of the endpoints).

##### 1.3.2 Simulations for the transient power law distribution

The probability density functions *g*(*T*) and *f*(*τ*) are given in Eqs. S3–24 and S3–26. Monte Carlo results are shown in Fig. S1–2 for *T*_1_ = 1, *T*_2_ = 125, *m* = 2.25. Again the Monte Carlo results agree well with the analytical solution at the scale of a log-log plot of the entire data set, and again a plot of the ratio is needed for a more stringent test. In Fig. S1–2 *b*, the ratio is in the ±1% band except for the noisy regions at small and large times.

Getting agreement this good, however, requires some modifications of the approach. The minor changes are increasing the number of mean escape times from 10^4^ to 6 × 10^4^, and extending the range of the histogram from [1/50, 2500] to [1/2500, 2500]. The major change is the use of importance sampling, which is the Monte Carlo equivalent of the standard practice of using a set of different frame rates in SPT experiments. The aim in both cases is to sample rare long-time events adequately. Normanno et al. (3) included measurements with continuous imaging, and time-lapse measurements with frame intervals of 100 ms, 500 ms, and 1 s [their Supplementary Note 7 and Supplementary Fig. 30]. In the Monte Carlo runs here, we divide the time interval *T*_1_ = 1 to *T*_2_ = 125 into four equal segments on a logarithmic scale, with boundaries 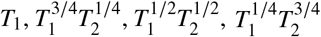, and *T*_2_, giving here 1, 3.344, 11.18, 37.38, and 125. An equal number of *T*-values are generated on each of the four intervals, and the resulting four histograms of *τ* are added with appropriate weighting. In the example here, the original area fractions of the four segments are 0.7807, 0.1727, 0.03819, 0.008446, and importance sampling changes the weighting to 0.25 each. So the fourth region is sampled almost 30× more than in simple sampling.

**Figure S1–2:**
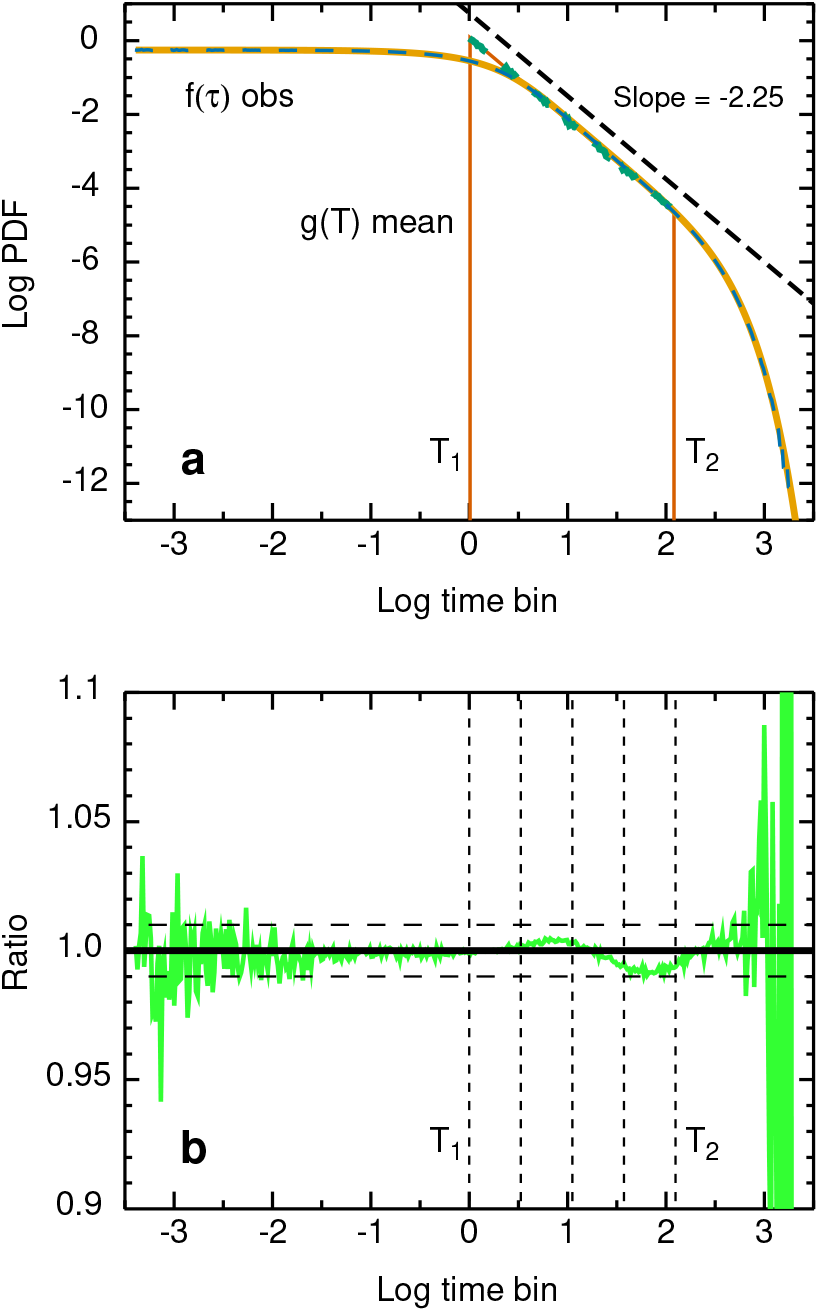
Results of Monte Carlo simulations in Mathematica for the TPL distribution *T*_1_ = 1, *T*_2_ = 125, *m* = 2.25. *(a)*, Log-log plot of PDFs versus time. Histograms of the TPL distribution of mean escape times *g*(*T*), analytical (red) and Monte Carlo (dashed blue-green), and the resulting observed distribution *f*(*τ*), analytical (orange) and Monte Carlo (dashed blue). Dashed line, slope −2.25. *(b)*, Ratio of Monte Carlo results to analytical results at the bin center (taken to be the geometric mean of the endpoints). Horizontal lines: ratio = 1 and the ±1% band. Vertical lines, boundaries for importance sampling. The corresponding ratio plot for the mean escape times *T* is not interesting enough to include. It shows good agreement, though it is noisy because the histogram for *T* is based on only 6 × 10^4^ data points, whereas the histogram for *τ* is based on 6 × 10^8^ data points. Note that the parameter in the Mathematica Pareto distribution differs from the exponent by 1.

The usefulness of importance sampling is shown in Fig. S1–3, for smaller-scale runs, 10^4^ values of *T* with 10^4^ values of *τ* for each *T*. Panel *(a)* shows the ratio for simple Monte Carlo, three independent repetitions, and panel *(b)* shows the ratio for 4-segment importance sampling, 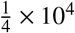 values of *T* in each segment with 10^4^ values of *τ* for each *T*, three independent repetitions. The vertical scales are the same. Importance sampling tames the noise significantly. In both cases, the three independent repetitions give clearly distinct curves, evidence that the ratio represents low-frequency noise, not systematic error.

**Figure S1–3:**
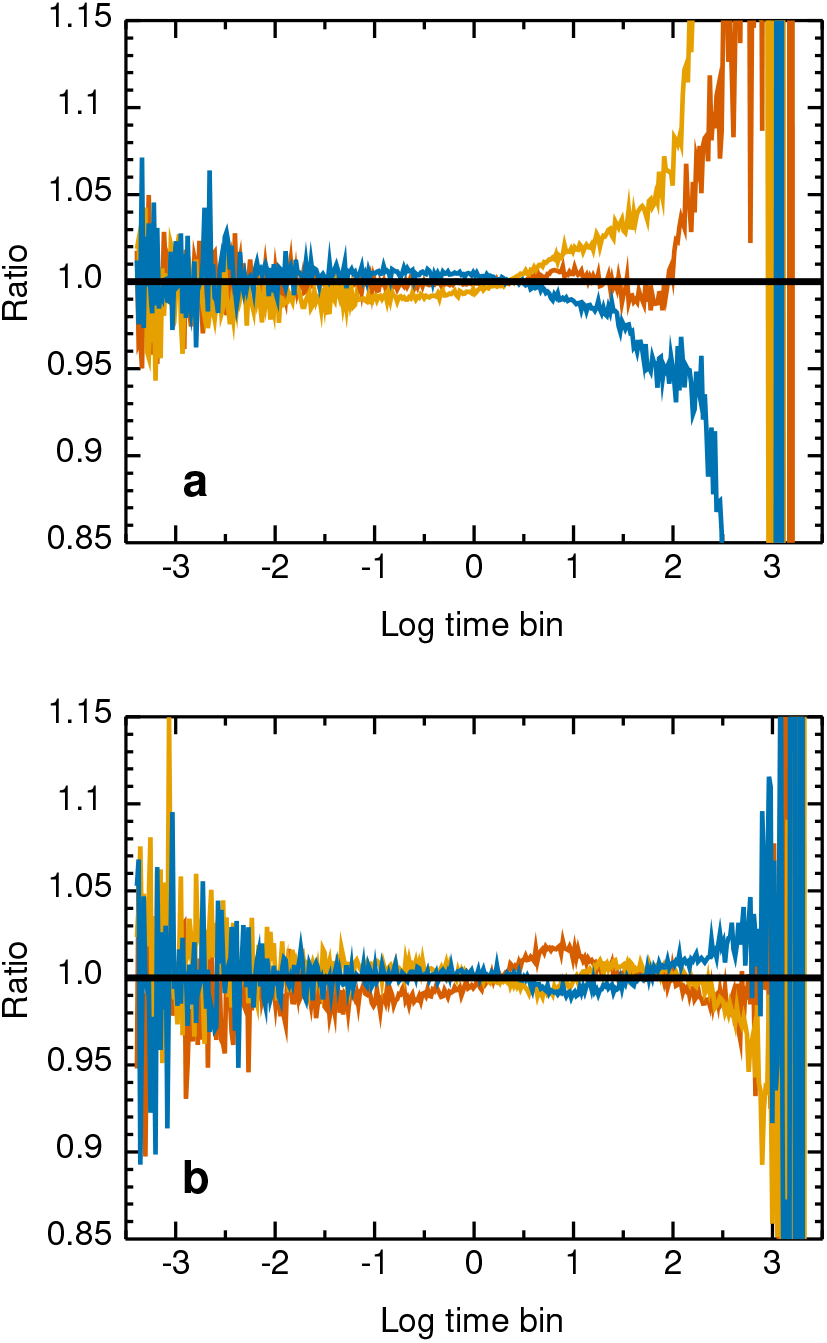
Ratio plots show the effect of importance sampling. Three independent runs for *(a)*, simple Monte Carlo, and *(b)*, Monte Carlo with 4-interval importance sampling. All six runs use the same total number of samples, 10^4^ values of *T* and 10^4^ values of *τ* for each *T*. For the TPL distribution *T*_1_ = 1, *T*_2_ = 125, *m* = 2.25, and trap fraction 1. These plots imply that the deviations from unity are low-frequency noise. Systematic error would be more systematic.

### 2 THE TRANSIENT POWER LAW DISTRIBUTION

#### 2.1 Experimental distributions

Table S2–1 summarizes the experimental TPL curves, which were presented in the original publications as plots of log *S*(*t*) versus log *t*, where *S*(*t*) is the survival function (text Eq. 2) and *m* is the corresponding exponent in text Eq. 3.

**Table S2–1:** Experimental results

| DNA-binding species | \|slope\| | $m$ | Time range | Ref. |
| --- | --- | --- | --- | --- |
| TetR | 0.69 | 1.69 | 200 ms to 70 s | Normanno et al. (3), Suppl. Fig. 30 |
| LacI | 1.12 | 2.12 | 30 ms to 10 s | Caccianini et al. (4), Fig. 3 |
| CRISPR-Cas 9 | $1.33 \pm 0.2$ | $2.33 \pm 0.2$ | 120 ms to 37 s | Knight et al. (5), Fig. 5-10 |

All these measurements were on mammalian cells, the CRISPR-Cas 9 experiments on NIH 3T3 mouse cells, and the others on U2OS human osteosarcoma cells. In all three papers, the final power-law histograms were constructed by combining several experimental histograms taken at different frame rates, as shown in the original figures. Knight et al. (5) gave the numerical value in terms of the inverse slope. All three measured distributions have a width of 2.5 decades, suggesting that the range might result from experimental constraints.

In my Monte Carlo calculations, the typical value of *m* is taken to be 2.25, the average of the values from Caccianini et al. (4), and from Knight et al. (5). A wide range of *m* is used to include the value from Normanno et al. (3), and the error bars of (5).

#### 2.2 Mathematical properties

Recall that the TPL probablility density function is

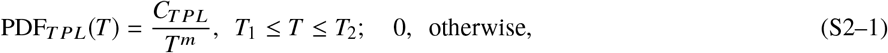

where *T* is a random variable, here either the mean or the observed escape time. Then for *m* ≠ 1

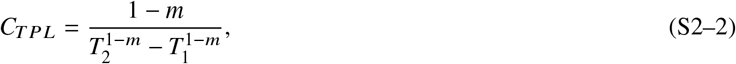

and the cumulative density function is

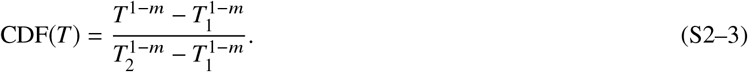

The CDF is important because it gives the survival function (text Eq. 2) and it is used to generate TPL random variates by the standard transformation method (18). If *r* is a random variate uniformly distributed on (0,1) and *x* is a random variate from the distribution described by a CDF, then take

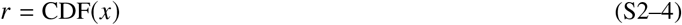

and invert to give

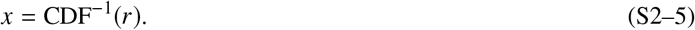

For the TPL this procedure gives

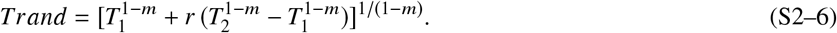

##### 2.2.1 Averages

The *n*th moment is

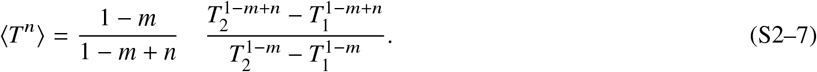

This expression holds for positive and negative *n*, so it also gives the moments of *W* = 1/*T*. The *n* = 0 case verifies the normalization. Note that 〈*T*〉 is indeterminate for *m* = 2 and must be evaluated by L’Hôpital’s rule to give 〈*T*〉 = *T*_1_*T*_2_(ln *T*_2_ – ln *T*_1_)/(*T*_2_ – *T*_1_), and similarly for the other cases in which *m* = *n* + 1. The median is

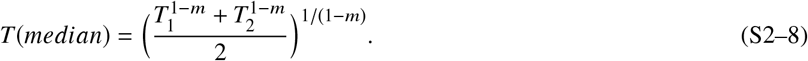

which is called the generalized mean (117) or the power mean (118).

If *m* = 1,

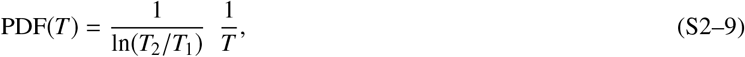

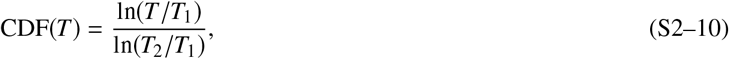

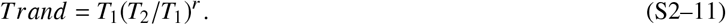

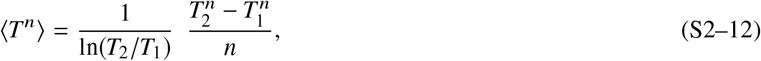

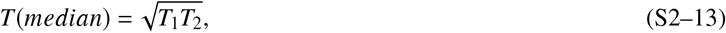

##### 2.2.2 Thermal averages

Thermal averages of moments of *T* and *W* are readily obtained to give the final state of the system. The Boltzmann factor for the TPL is (text Eq. 8) *P_i_* = *T_i_*/∑_*i*_ *T_i_*, so the thermal average nth moment of *T* is

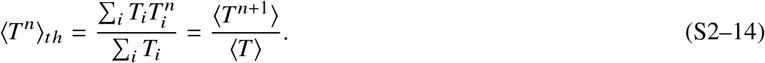

This expression holds for positive and negative n, so it also gives the moments of *W* = 1/*T*. The *n* = 0 case verifies the normalization.

##### 2.2.3 Binding energies

The binding energy is

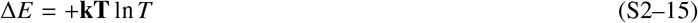

so the mean binding energy is, in units of **kT**,

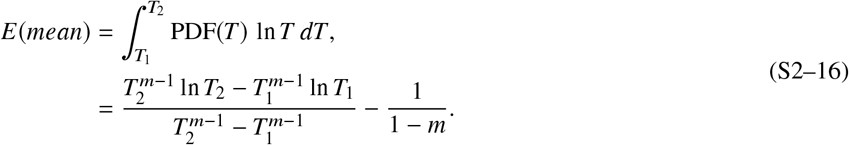

on integration by parts.

### 3 FROM BINDING ENERGY DISTRIBUTIONS TO OBSERVED ESCAPE TIME DISTRIBUTIONS

Recall from the text that in this section it is necessary to distinguish between the mean escape time T and the observed escape time *τ*.

#### 3.1 Truncated binding energy distributions to mean escape time distributions

In the text we showed the standard result that escape from a pure exponential distribution of mean binding energies according to an Arrhenius factor yields a power-law distribution of mean escape times. Here we give the results for three truncated binding energy distributions: exponential, Gaussian, and log-normal. The full expressions are complicated due to the truncation, but for each binding energy distribution, the effect of truncation can be expressed as a common normalization constant for *ϕ*(Δ*E*) and *g*(*T*).

##### 3.1.1 Truncated exponential

A truncated exponential distribution of binding energies yields a truncated power-law distribution of mean escape times. The normalized PDF for a truncated exponential distribution is

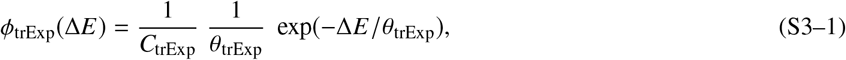

where

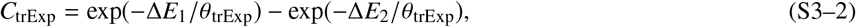

Δ*E*_1_ < Δ*E*_2_ are the lower and upper limits, and *θ*_trExp_ is a width parameter in units of energy, not related in a simple way to the mean or median energy. We use text Eq. 12 to obtain

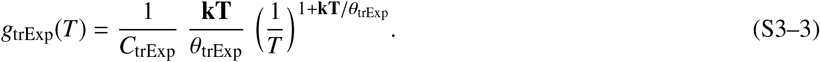

This equation is the product of the pure power law distribution Eq. 14, the normalization factor Eq. S3–2 for truncation, and the ratio of the width parameters. In the limit Δ*E*_1_ → 0, Δ*E*_2_ → ∞, *C*_trExP_ → 1.

##### 3.1.2 Truncated Gaussian

A truncated Gaussian distribution of binding energies yields a distribution of mean escape times clearly distinct from a power law. The normalized PDF for a truncated Gaussian distribution is

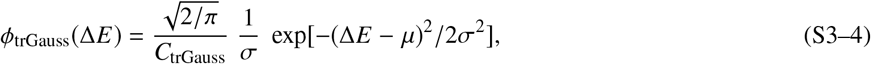

where

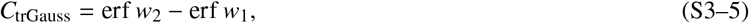

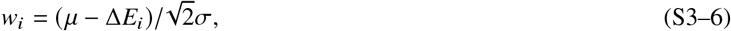

erf *w* is the error function, *μ* is the mean of the Gaussian distribution, *σ* is its standard deviation, and the other quantities are defined as above. In the limit Δ*E*_1_ → 0, Δ*E*_2_ → ∞, *f*_trGauss_(Δ*E*) reduces to the expression for a Gaussian restricted to positive Δ*E*

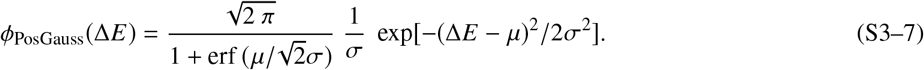

Proceeding as before, we obtain from text Eq. 12

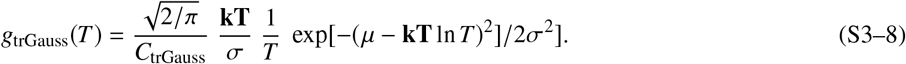

Here **kT**, *μ*, and *σ* have dimensions of energy, and T is dimensionless.

##### 3.1.3 Truncated log-normal

A truncated log-normal distribution of binding energies yields approximately power-law mean escape times over a limited but significant range. In one parameterization, the PDF for a nontruncated log-normal distribution is

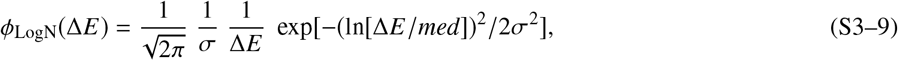

where *med* is the median of Δ*E*, and the shape parameter *σ* is the standard deviation of ln[Δ*E*/*med*]. Zaninetti (119) has a useful collection of formulas for this parameterization. The normalized PDF for a truncated log-normal distribution is

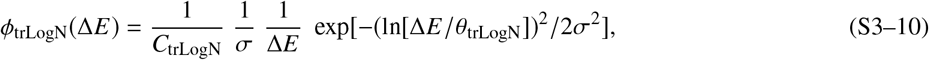

where

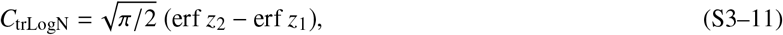

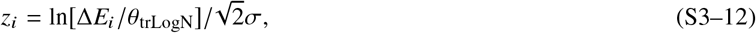

and *θ*_trLogN_ is a width parameter. In the limit Δ*E*_1_ → 0, Δ*E*_2_ → ∞, Eq. S3–10 reduces to Eq. S3–9. Proceeding as before, we obtain

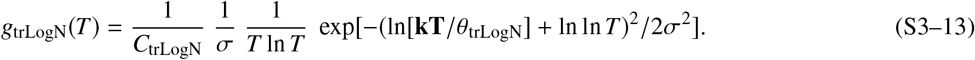

In Eq. S3–13, *C*_trLogN_, *T*, and *σ* are dimensionless; Δ*E*, θ_trLogN_, and **kT** have dimensions of energy.

Why does this PDF give an approximately log-log form? Rearrange S3–13 and collect the various constants into coefficients *A_i_* to give

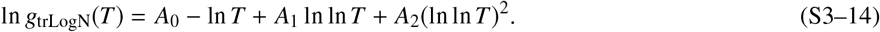

So *g*_trLogN_(*T*) is not really a power-law distribution, though the tail can approximate one over a significant range.

We examine the dependence on **kT** numerically. Fig. S3–1 shows plots of log *g*_trLogN_ versus log *T* for *μ* = 1, *σ* = 1, truncation at Δ*E*_1_ = 1, Δ*E*_2_ = 6.75. The truncation is chosen to give a distribution of mean escape times of width 2.5 decades as in the Dahan experiments. The values of **kT** are 1/2, 1, 3/2, and 2. Truncation makes the distribution approximately power-law, with the power tunable to some extent through **kT**, but major deviations from linearity occur if the limits include the peak of the log-normal distribution of Δ*E*.

**Figure S3–1:**
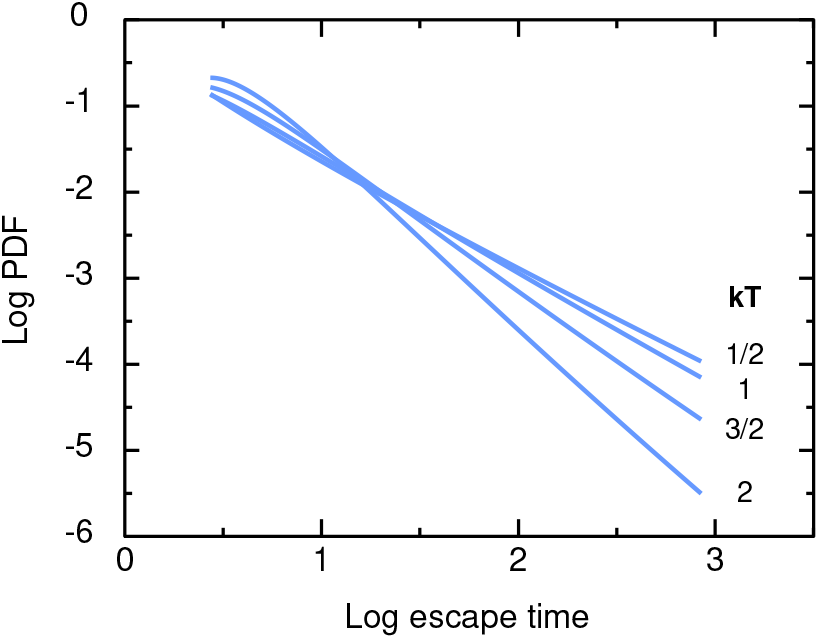
Mean escape time distributions for a truncated log-normal distribution of binding energies. Plot of log *g*_trLogN_ from Eq. S3–13 versus log *T* for the indicated values of **kT**, with *μ* = 1, *σ* = 1, Δ*E*_1_ = 1, Δ*E*_2_ = 6.75.

#### 3.2 Mean escape time distributions to observed escape time distributions

As stated in the text, there are two distinct escape times here, the observed escape time *τ*, and the mean escape time *T*, which is determined by the binding energy in the case of escape from a binding site, and by the obstacle geometry and dynamics in the case of escape from a geometrical trap. The distribution of *τ* is broader than the distribution of *T* due to the scatter in *τ* for a fixed value of *T*.

If a tracer escapes from a trap with mean escape time *T* according to the Arrhenius law, the observed escape time *τ* for each visit is a random number from an exponential distribution

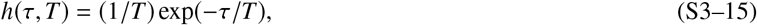

or the analogous discrete form, the geometrical distribution, in the Monte Carlo simulations. The resulting probability distribution *f* for the observed escape times *τ* is a compound distribution *f*(*τ*) in which the parameter *T* is distributed according to a PDF *g*(*T*), giving in general

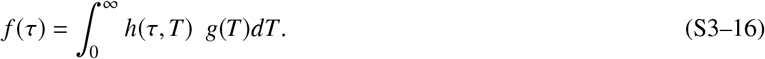

All the functions are PDFs so each one must be normalized individually. Here

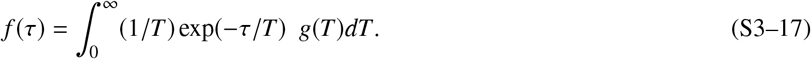

The problem is therefore a Fredholm integral equation of the first kind: Given an observed *f*(*τ*), find the underlying distribution *g*(*T*) of mean escape times.

These integral equations are common in a variety of contexts, such as anomalous diffusion (120), the distribution of activation energy barriers for a two-state reaction (121), kinetics of peptide folding (122), light scattering by polydisperse particles (123), luminescence decay (124), NMR signals in terms of the distribution density of NMR relaxation times (NMR relaxometry) (125), and recombination kinetics after photodissociation of CO-myoglobin (126). These integral equations are well-known to be ill-posed; numerical inversion is highly sensitive to noise in the observed *f*(*τ*).

Here we adapt the derivation of Metzler et al. (122), one of the shortest and clearest treatments of the problem. We change the variable in the integral equation S3–17, with *s* = 1/*T, dT* = −*ds*/*s*^2^, to obtain a Laplace transformation 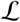

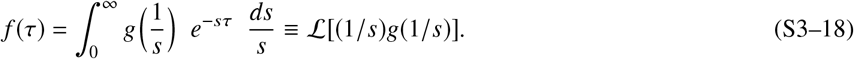

Then

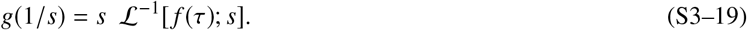

Finally, return to the original variable

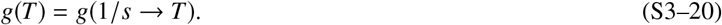

The integral equation thus reduces to an inverse Laplace transformation. The problem is still ill-posed – the inversion involves an integral over the positive exponential exp(+*sτ*) so noise is amplified – but the inversion is important enough that the problem has received much attention, including two outstanding reviews (127, 128).

The direct solution would start with the experimental PDF *f*(*τ*) of the observed escape times and invert this according to Eq. S3–19 to give the PDF *g*(*T*) of the mean escape times. But this inversion is problematic, both analytically and numerically, so we use an indirect approach. Assume that the mean escape time *T* follows a TPL distribution, a power law on grounds of the experimental power law, truncated on physical grounds. Then generate the PDF of the observed times *τ* from this.

We find the analytical solution as follows. First, consider the nontruncated case, and ignore the requirements that a PDF be bounded and normalized. Then the analytical solution is trivial. If

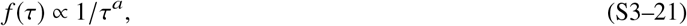

then

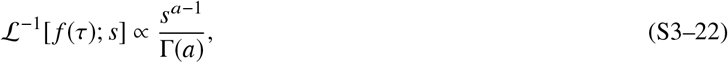

where Γ(*a*) is the gamma function, and

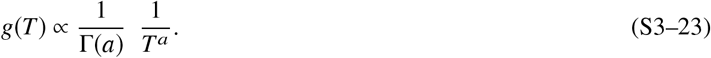

A power-law distribution of observed escape times *f*(*τ*) implies a power-law distribution of mean escape times *g*(*T*). The complications arise from the truncation. We assume that *g*(*T*) is a TPL distribution (text Eqs. 3,4),

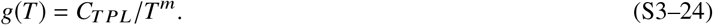

and substitute this into Eq. S3–17, putting in the truncation, to give

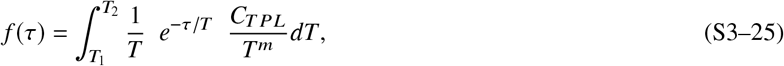

which gives by a simple change in variable *x* = *τ*/*T*

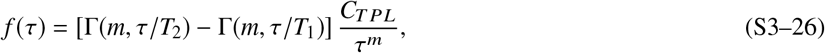

where Γ(*m, z*) is the incomplete gamma function

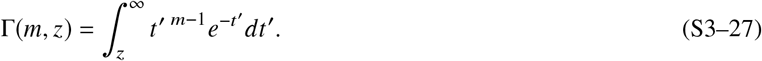

The exponents are the same in *g* and *f* on account of the normalization factor 1/*T* in *h*. The form is simple. The power law is the same as that obtained in the nontruncated case, but it is multiplied by a truncation function

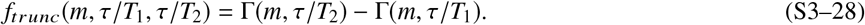

The behavior of *f_trunc_* can be understood from the properties of Γ(*m, τ*/*T*). As *τ*/*T* → 0, Γ(*m, τ*/*T*) → Γ(*m*). The corner of the gamma function occurs at the maximum in the derivative with respect to *τ*/*T*, at *τ*/*T* = *m* − 1. The lower cutoff is determined by the fact that *f_trunc_* is the difference of two Γ functions with the same *m*. The Γ functions mostly cancel out at small *τ*/*T*, and the flat region in *f_trunc_* begins when Γ(*m, τ*/*T*_1_) is negligible, here around *τ*/*T*_1_ = 5. When *T*_1_ << *T*_2_ as here, the corner in *f_trunc_* coincides with the corner in Γ(*m, τ*/*T*_2_) and occurs at log *τ* = log *T*_2_ + log(*m* − 1). For *m* = 2.25, log(*m* − 1) = 0.0969 so the corner is close to log *T*_2_.

Fig. S3–2 shows the PDFs *f*(*τ*) and *g*(*T*) for a TPL distribution with *T*_1_ = 1, *T*_2_ = 200, and *m* = 2.25. (Note that these times are defined on [0, ∞), that is, they are the additional delay introduced by the traps. In contrast, the times in the Fortran Monte Carlo simulations are defined on [1, ∞) because a move there always requires at least one time step.) Panel *(a)* is a linear plot and panel *(b)* is a log-log plot. The linear plot shows that most of the probability density is at short times. The log-log plot shows the power-law structure: for the mean escape times *T*, a truncated power law, and for the observed times *τ*, a truncated power law modified at short and long times by the escape statistics. Panel *(c)* shows the form of *f*(*τ*): the product of a power-law distribution and a truncation function. The truncation function is flat in a region between *T*_1_ and *T*_2_. For small *t* with *T*_1_ << *T*_2_, a series expansion shows that the truncation function is proportional to *τ^m^*, canceling the power-law factor so that *f*(*τ*) is indeed flat for small *τ*. Panel *(d)* shows the outcome: *f*(*τ*) is flat at small times, and the extent of the power-law region increases as *T*_2_ increases. Also included is the PDF for the nearly delta-function case *T*_1_ = 1, *T*_2_ = 1.001, *m* = 2.25, to show the shape of the nearly pure exponential PDF Eq. 13 in a log-log plot. All the PDFs have similar shapes, flat at small times and steeply decreasing at large, but the pure exponential case has no power-law region. Corresponding Monte Carlo results are presented in Suppl. 1.3.

In summary, a TPL distribution *g*(*T*) of mean escape times *T* yields a mathematically well-defined broadened distribution *f*(*τ*) of observed escape times *τ* as shown in panel *(b)*. But experimental measurements will likely yield a truncated version of this *f*(*τ*), truncated at short times due to the definition of a binding event, and truncated at long times because the finite number of observations limits the number of rare events detected. Both truncations are dependent on details of the experiment and data analysis.

**Figure S3–2:**
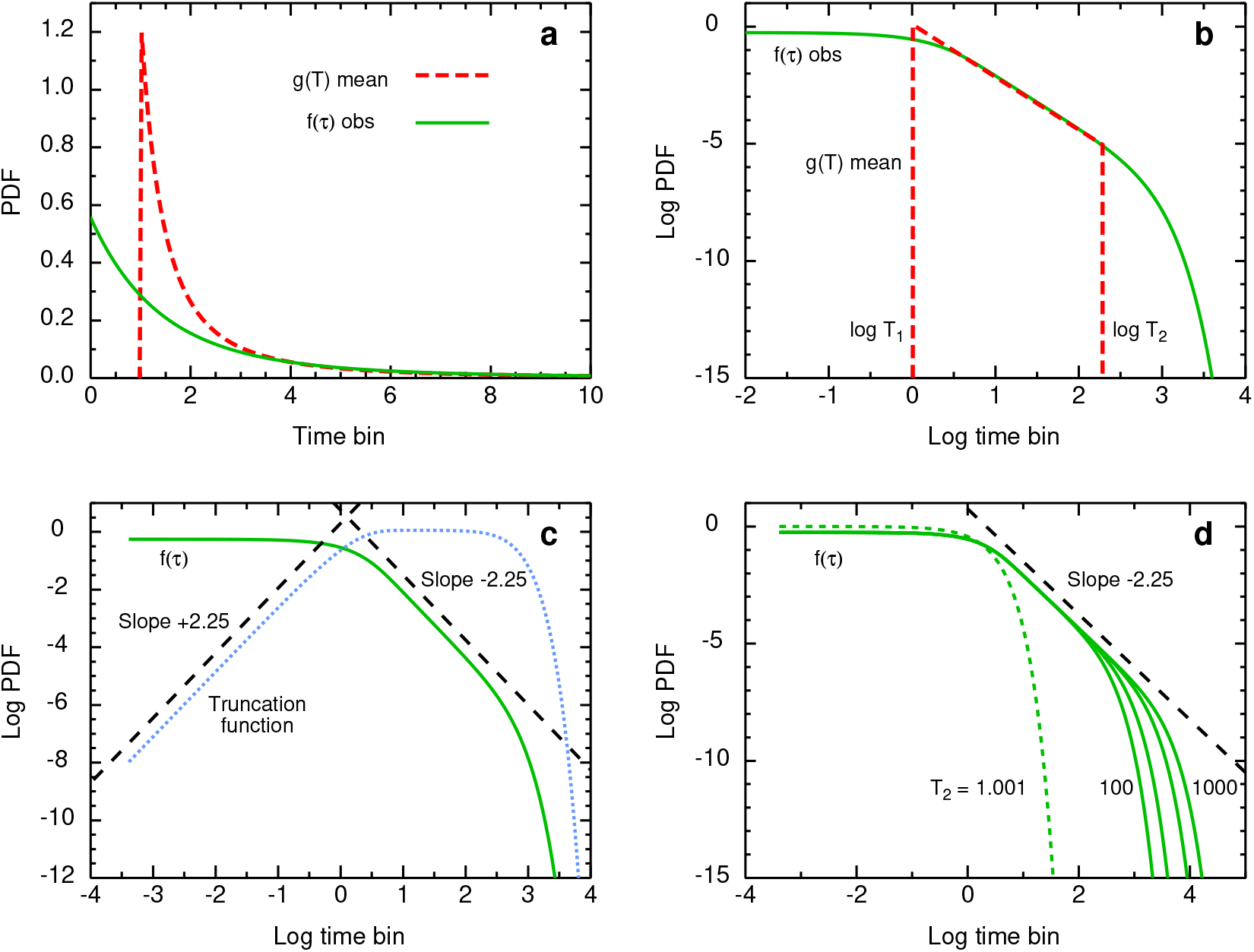
Probability density functions *f*(*τ*) (solid green) and *g*(*T*) (dashed red) for TPL distribution with *T*_1_ = 1, *T*_2_ = 200, and *m* = 2.25. *(a)*, Lin-lin plot of *f*(*τ*) and *g*(*T*). **(b)**, Log-log plot of *f*(*τ*) and *g*(*T*). *(c)*, Log-log plot of *f*(*τ*) and truncation function *f_trunc_*(*τ*) (dotted blue). Dashed lines, slopes ±2.25. *(d)*, Log-log plot of *f*(*τ*) for *T*_1_ = 1, *T*_2_ = 100,200,500,1000, *m* = 2.25. Dashed line, slope = −2.25. As *T*_2_ increases, the width of the power-law region in *f*(*τ*) increases. For comparison, the dotted line shows the case *T*_2_ = 1.001 to approximate a pure exponential *f*(*τ*) resulting from a delta-function *g*(*T*).

### 4 PLOTS OF LOG D(T) VERSUS LOG T

#### 4.1 Analysis

Text Fig. 6 showed families of log *D*(*t*) versus log *t* curves as one of the TPL parameters was varied – *T*_1_, *T*_2_, *m*, and trap concentration – with the others held constant. These plots give a qualitative picture of how the TASD parameters depend on the TPL parameters, For a more quantitative picture, we analyze the curves as in Suppl. 1.2 and plot the resulting TASD parameters in Fig. S4–1. Here *m, T*_1_, and *T*_2_ are considered.

When *T*_2_ is varied with *T*_1_ = 1 constant (panel *a*), diffusion is mildly anomalous for the standard case *m* = 2.25 and somewhat more anomalous for *m* = 0.30, a value chosen to match earlier work on discrete trap hierarchies (Suppl. 4.2). Diffusion is less anomalous than for an obstructed random walk at the percolation threshold for the simple cubic lattice (129). The log-log plot of *t*(*cross*) versus *T*_2_ is approximately linear (panel *d*), with slope around 2/3 for both values of *m*. When *T*_1_ is increased at constant *m* = 2.25 and *T*_2_ = 200, diffusion becomes more normal (panel *b*) because the range of trap depths decreases. The log-log plot of *t*(*cross*) versus *T*_1_ is approximately linear, with slope around 1/3 (panel *e*). When *m* is increased at fixed *T*_1_ and *T*_2_, *α* shows a minimum and *t*(*cross*) shows a maximum, at different values of *m*. That is, for fixed *T*_1_ and *T*_2_, there is a “most anomalous” and a “longest anomalous” case. These extrema are presumably related to the extrema in *D*(0) – *D*(∞) and *D*(∞)/*D*(0) as m is varied (text Eqs. 19, 23), and will be examined in detail in future work. Panels *(e)* and *(f)* explain the mildness of the TASD in panels *(a)* and *(c)*. Neither *m* = 0.30 nor *m* = 2.25 is near either extreme case.

The dependences are complicated, and will provide a stringent test of future theoretical analysis of the model. The dependence on trap fraction is better described in terms of rescaling, as described in section 4.2.

**Figure S4–1:**
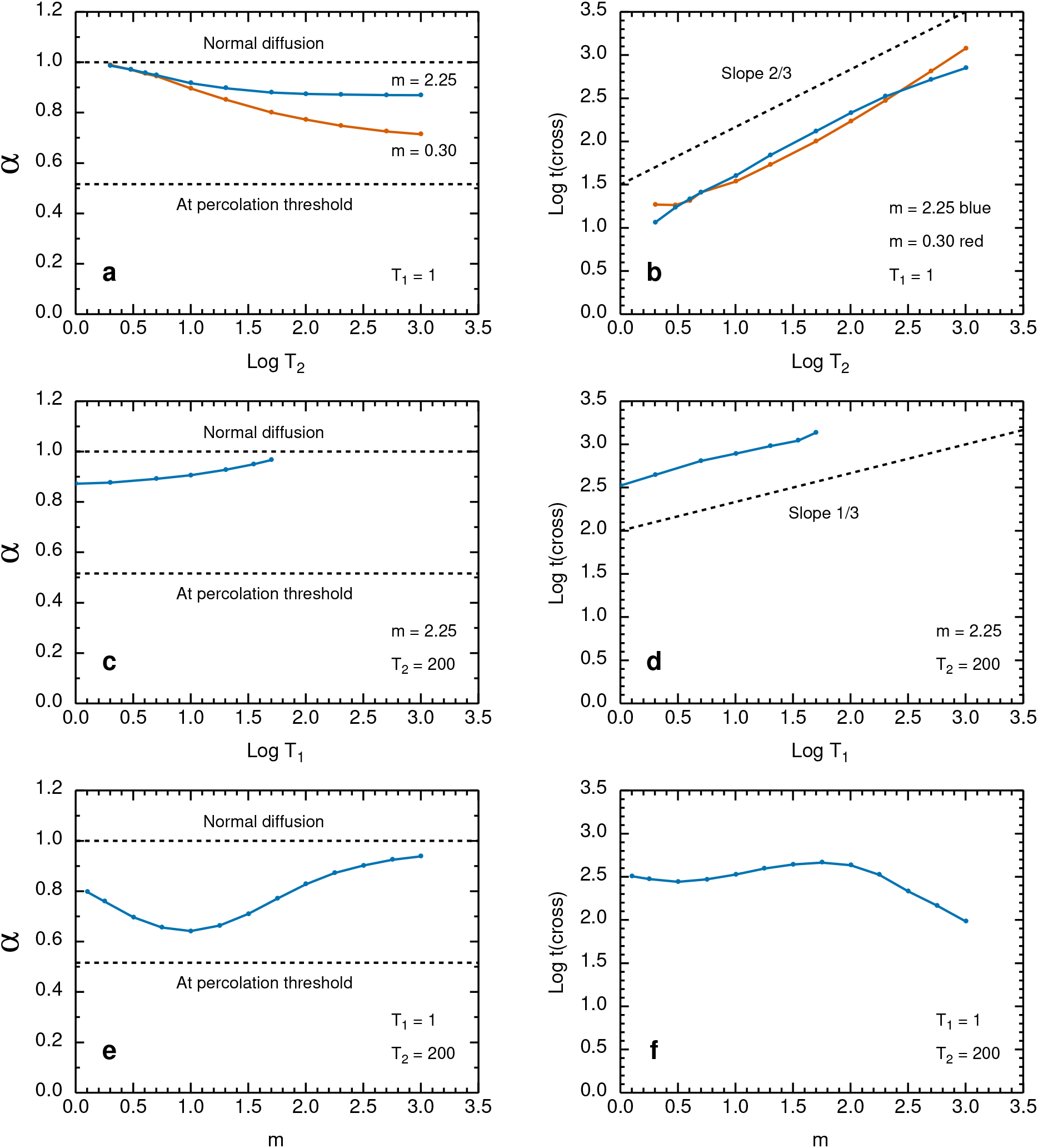
Variation in the TASD parameters *α* (left column) and log *t*(*cross*) (right column) as a function of one TPL parameter at constant values of the other TPL parameters. The panels are shown with constant axes for *α* and for *t*(*cross*) to make clear the differences in sensitivity of the results to the TPL parameters. Panels *(a)* and *(b)*. Vary *T*_2_ at constant *T*_1_ = 1, *m* = 0.30 and 2.25, trap fraction 1. The dashed line in *(b)* has slope 2/3. Panels *(c)* and *(d)*. Vary *T*_1_ at constant *T*_2_ = 200, *m* = 2.25, trap fraction 1. The dashed line in *(d)* has slope 1/3. Panels *(e)* and *(f)*. Vary m at constant *T*_1_ = 1, *T*_2_ = 200, trap fraction 1.

Earlier work examined TASD due to a discrete hierarchy of traps (1), described in terms of the recipe of Shlesinger (130) for fractal time: a hierarchy of traps with an order of magnitude longer escape times, an order of magnitude less often, where the orders of magnitude are not necessarily in base 10 or in the same base. This work led to a simple qualitative summary: As the hierarchy of traps deepens, TASD is more anomalous for a longer time. Why is the qualitative picture more complicated in the current work than in the earlier work? One reason is that we examine a wider range of parameters here. Another is that we examine the results more quantitatively, extracting *t*(*cross*) and *α*, and comparing curve shapes by rescaling (Suppl. 4.2). There are also differences in the models: a 2D triangular lattice in the old versus a 3D simple cubic in the new, though both have coordination number 6; discrete levels of traps in the old versus a continuum in the new, though both are based on a power-law dependence.

#### 4.2 Rescaling of log D(t)

The trap concentration is a major factor in determining *α*, but text Eqs. 19 and 23 for *D*(0) and *D*(∞) account for much of the concentration dependence. If we rescale plots of log *D*(*t*) linearly so that the dependent variable is

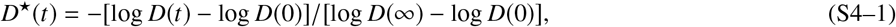

the range of slopes is decreased. Three examples are shown in Fig. S4–2, the unscaled Monte Carlo data in the left column and the rescaled in the right. This normalization by *D*(0) and *D*(∞) will be an essential step in future work on the full scaling analysis of the Monte Carlo data.

The standard case *T*_1_ = 1, *T*_2_ = 200, *m* = 2.25 shows almost complete data collapse as the trap concentration is varied, as shown in Fig. S4–2, panels *(a)* and *(b)*. These are essentially one curve, stretched vertically according to the values of *D*(0) and *D*(∞). Other parameter values give much less data collapse, panels *(c)* and *(d)*. Changing m from 2.25 to 0.30 with *T*_1_ and *T*_2_ unchanged gives a family of clearly distinct curves on rescaling (data not shown).

We use this rescaling to compare current TPL results with earlier work on TASD due to discrete hierarchies of traps (2), where a typical example had 16 trap sites with escape time *T*, 8 with *T*^2^, 4 with *T*^3^, and 2 with *T*^4^, say with *T* = 10. The occupancy decreases by a factor of 2 at each level, giving a discrete power-law distribution with exponent *m* = log 2 = 0.3010. The value of *T*_1_ is 1 in both discrete and TPL cases, and *T*_2_ was chosen to be 2000 so that the values of *D*(∞) for a trap fraction around 0.5 were similar in the discrete and TPL cases. Results for the TPL with *m* = 0.30, *T*_1_ = 1, and *T*_2_ = 2000 are shown in panels *(e)* and *(f)*. Rescaling reduces the variation in the family of curves for the different concentrations, but not to the extent of the *m* = 2.25 case. The corresponding plots for discrete traps (2) are shown in panels *(c)* and *(f)*. Here *D*(0) and *D*(∞) are from text Eqs. 19, 23 as in the TPL case but the averages are for the appropriate distribution of discrete traps. The extent of data collapse is similar in the continuum and discrete cases, but the shapes of the curves are different. For example, the values of *t*(*cross*) are around a factor of 10 greater in the discrete case.

**Figure S4–2:**
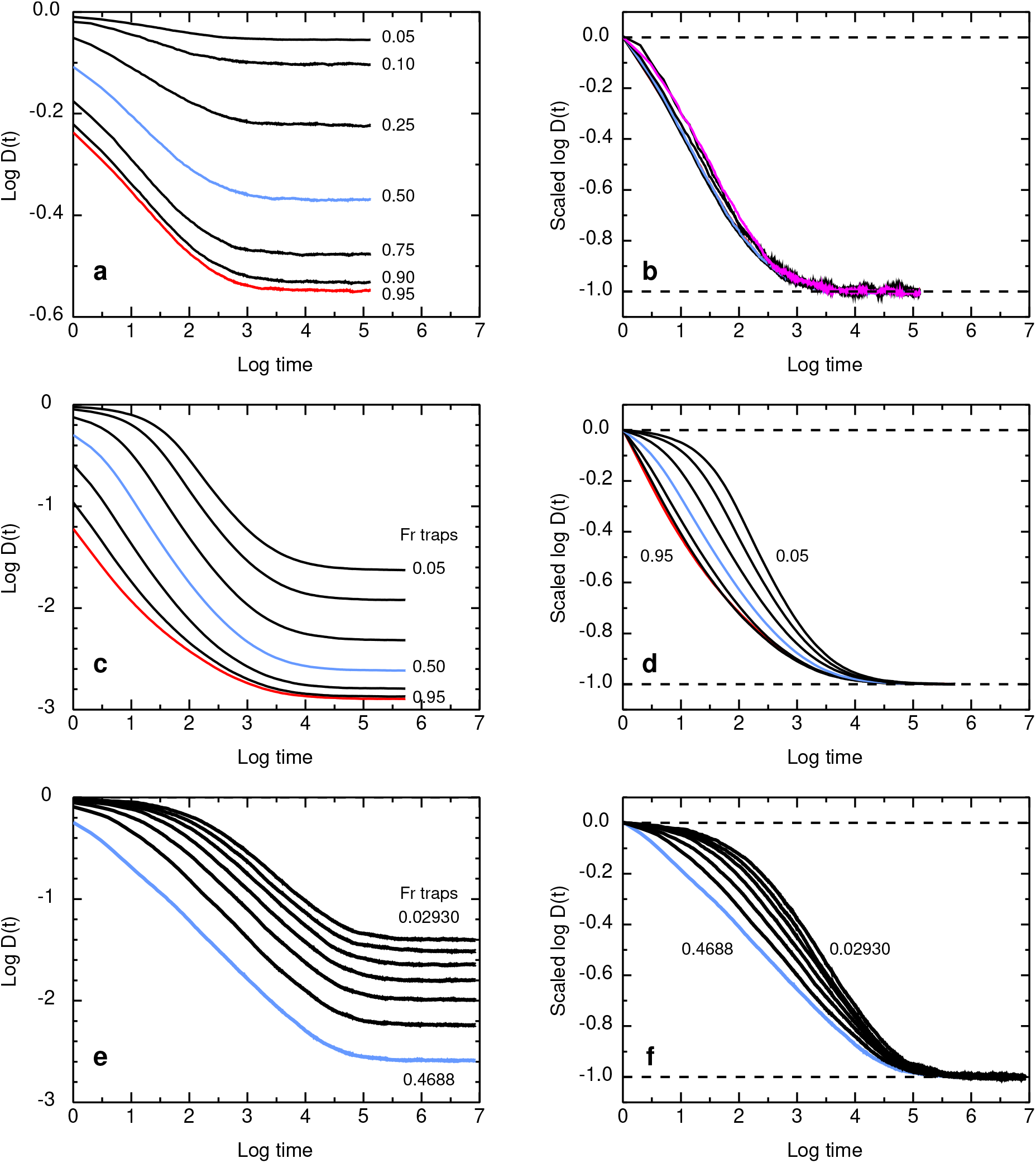
Extent of data collapse on rescaling. Log-log plots of *D*(*t*) versus *t* for various trap concentrations. Left column, unscaled Monte Carlo data. Right column, the Monte Carlo data rescaled according to Eq. S4–1. Panels *(a)* and *(b)*. The standard test case *T*_1_ = 1, *T*_2_ = 200, *m* = 2.25, with trap fractions 0.05, 0.10, 0.25, 0.50 (blue), 0.75, 0.90, and 0.95 (red). Panels *(c)* and *(d)*. The same for *T*_1_ = 1 but with *T*_2_ = 2000 and *m* = 0.30 to approximately match D(∞) for the discrete case for trap fractions around 0.5. Panels *(e)* and *(f)*. Earlier results for discrete levels of traps on the triangular lattice (Fig. 4 *c* of Ref. (2)). There are 16 traps with mean escape time 10, 8 with time 10^2^, 4 with time 10^3^, and 2 with time 10^4^. Trap concentrations given are for a single set of 30 traps on lattices with edges 32, 28, 24, 20, 16, 12, and 8, giving trap fractions of 0.02930, 0.03827, 0,05208, 0.07500, 0.1172, 0.2083, and 0.4688.

#### 4.3 How much averaging is needed to see the structure?

The plots of Monte Carlo data for log *D*(*t*) versus log *t* show very little noise; the simulations are typically run for 500 sets of traps, and 2000 tracers for each set of traps. Results are ensemble-averaged; in systems showing aging, averaging within individual trajectories leads to entirely different behavior (24). In computer simulations it is easy to run 10^6^ trajectories to get good statistics, but in experiments on live cells, observing so many trajectories is difficult. How many repetitions are required for the ensemble-averaged curves to approach their final form? Fig. S4–3 shows a simulation addressing that question. Here a single trap configuration is generated, and log *D*(*t*) curves are generated averaging over 10, 20, 50, and 2000 tracers. Averaging over 50 tracers seems minimally appropriate.

**Figure S4-3:**
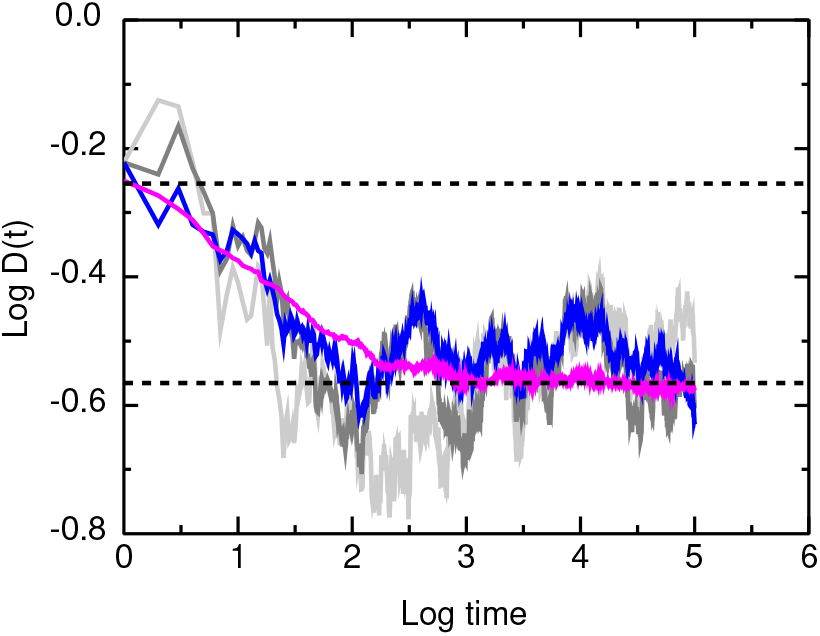
Plots of log *D*(*t*) versus log *t* for a single trap configuration, with *D*(*t*) averaged over 10 (light gray), 20 (dark gray), 50 (blue), and 2000 (magenta) independent trajectories.

#### 4.4 Effect of the initial delay in the measurements

Normanno et al. (3) measured tracer motion over a range of times 0.01 to 600 sec, with a delay time of 1 min between mixing and the start of observations. What effect does that delay have on the interpretation of the results in terms of a TASD model?

##### 4.4.1 Fill the time delay in various ways

One approach is to construct a system with a prescribed initial gap in the delay times, fill the gap in various ways, and compare the resulting plots of log *D*(*t*) versus log *t*. We start with a TPL distribution of mean escape times with *T*_1_ = 10, *T*_2_ = 200, *m* = 2.25. First, we fill the gap by extending the original power law to the smallest escape time, giving the TPL *T*_1_ = 1, *T*_2_ = 200, *m* = 2.25. Second, we keep the original power law and assume for all shorter times a random escape time *T* uniformly distributed on [1, 10], with the fractions matched at *T* = 10. Third, we keep the original power law and assume that for all shorter times *T* = 1, with the same fraction as in the previous case. Fig. S4–4 *(a)* shows these distributions as log-log plots, and Fig. S4–4 *(b)* shows the resulting TASD curves. The choice of distributions in the gap affects the range of *D*(*t*) considerably and affects the crossover time significantly, but none of the choices abrogates TASD. Clearly, measurements with shorter delays would be useful.

Aside: It is well-known that pure power-law distributions are scale-free, implying that a short observation window is sufficient to characterize the distribution. If we knew a priori that the distribution was power-law, we could extend the distribution outside the range of measurement. Unfortunately, the question here is, over what range is the distribution a power law?

##### 4.4.2 3D diffusion during the delay time

For simple 3D diffusion, a tracer can travel a significant fraction of the size of the nucleus in a one-minute initial delay. But the relevant quantity is not the distance but the number of distinct sites visited, and the tracer does not visit a significant fraction of the sites in that time. This fact is simply illustrated by Monte Carlo calculations for a random walk on a cubic lattice in a cube with reflecting boundaries. We compare the time dependence of the normalized mean-square displacement *f* (RSQ) = 〈*r*^2^(*t*)〉/〈*r*^2^(∞)〉 with the fraction *f* (DSV) of distinct sites visited. Results are shown in Fig. S4–5 for cubes of sides *L* = 30, 62, and 94. The *f* (RSQ) curves approach their limit at least a factor of 10 faster than the *f* (DSV) curves do, and the difference becomes greater for larger cubes.

**Figure S4–4:**
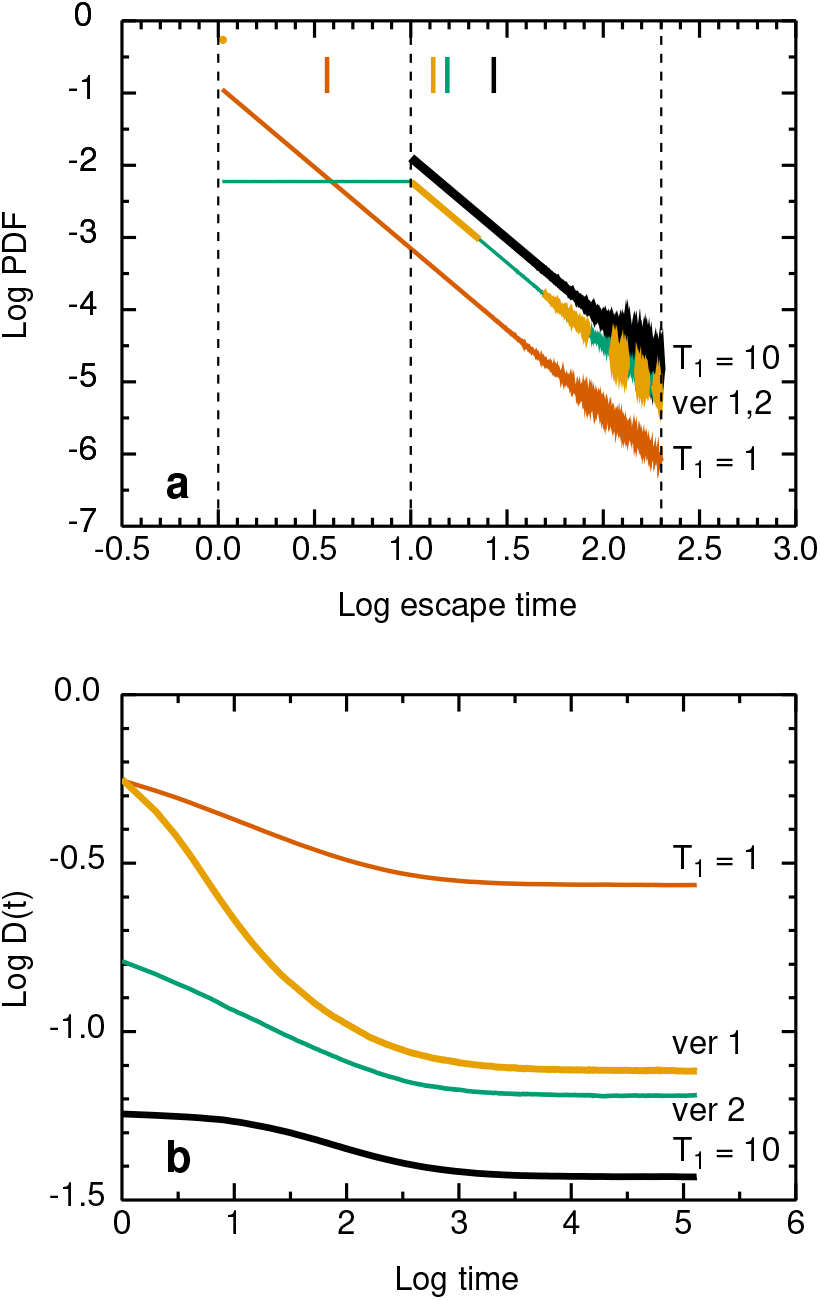
Effect of the initial gap in observation times on TASD. In all cases, *T*_2_ = 200, *m* = 2.25. *(a)*, Histograms of *T* as log-log plots, for the initial gap case *T*_1_ = 10, and three ways of filling the gap *T* = [1,10]. First, labeled *T*_1_ = 1, direct extension using the same TPL, that is, the TPL with *T*_1_ = 1. Second, labeled *ver 1*, filling the gap with random T uniformly distributed on [1,10]. The fraction of uniformly distributed sites is chosen to be 0.535368 so that the values of the uniform and TPL distributions match at *T* = 10. Third, labeled *ver 2*, the same fraction of *T* but all set to 1. *(b)*, The resulting plots of log *D*(*t*) versus log *t*, showing that the different choices change the TASD parameters significantly but do not abrogate TASD.

More quantitatively, we can find the crossover time in the *f* (RSQ) curve and obtain the fraction of distinct sites visited at the crossover time. The fractions are small: as the cube edge increases from *L* = 30 to 62 to 94, *f* (DSV) decreases from 0.0378 to 0.0192 to 0.0118. The size of the cube edge for a full-size model nucleus is significantly larger. If we take the lattice constant to be the experimental spatial resolution, we find *L* = (diameter of nucleus)/(resolution element) = 10 *μ*m/25 nm = 400, and if we take the lattice constant to be the size of a binding site, *L* becomes even larger.

Fig. S4–5 is simply one way of quantifying the classic argument for facilitated diffusion of Berg et al. (131) that a search of the nucleus by pure 3D diffusion is far slower than observed in a living cell, and a combined 3D/1D search is involved. The speedup of sampling by combined 3D/1D diffusion is discussed by Halford and Marko (132), and by Slutsky and Mirny (29), for example.

Technical aside: At the resolution of this figure, the curves look very similar. In fact, the curves for *f* (RSQ) and *f* (DSV) are superimposable if they are shifted so that their 50% points are at the same time. But this agreement is not exact. In the figure, f (DSV) is the ensemble average for a fixed time scale, with coarser resolution at large times (Suppl. 1.1). A rigorous simulation of *f* (DSV) to determine the dependence of the cover time on system size would be done differently. For each repetition, one would record the exact time step at which the actual last site is visited, with no artificial limit on the maximum time. and find the statistics from these exact values. A full treatment shows that the 3D cover time is proportional to *N* log *N*, where *N* is the number of lattice sites (133, 134). These limitations on the Monte Carlo results do not affect the argument that a tracer can diffuse a significant distance without visiting a significant fraction of lattice sites.

**Figure S4–5:**
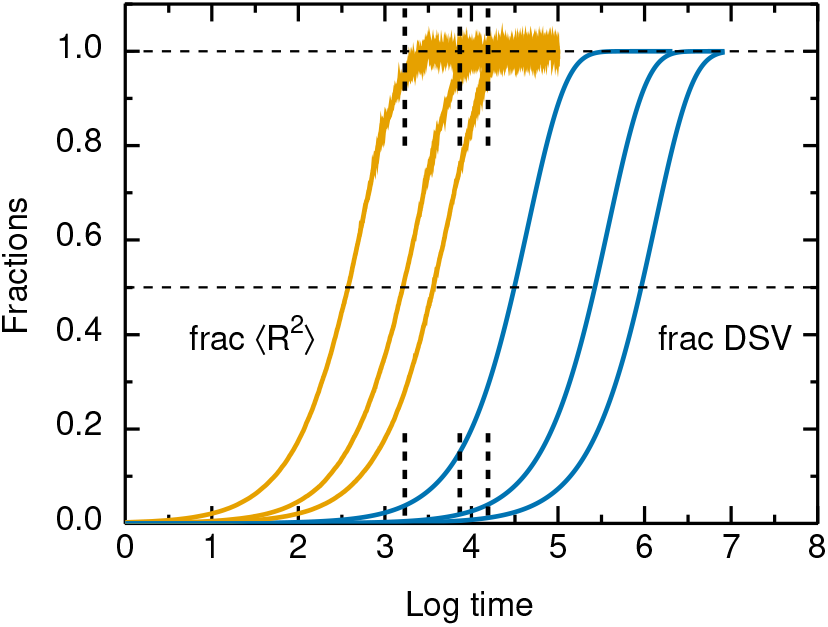
Mean fractional mean-square displacement and distinct sites visited as a function of time for random walks within a cube of edge *L* with reflecting boundaries. Plots of *f* (RSQ) (left 3 curves) and *f* (DSV) (right 3 curves) versus log *t* for *L* = 30, 62, and 94 (left to right). Vertical lines, the crossover time for *f* (RSQ), obtained by fitting 301 points around the 50% point to a straight line and extrapolating to 1.

##### 4.4.3 The moral of the story

First, it would be highly useful to reduce the delay time so we would have experimental facts, not just results of various models. Photoactivated probes would be useful, as discussed in the text. Second, Normanno et al. (3) said that it would be to know the concentration of binding sites in the nucleus for the molecule of interest. A good point, and a more specific form of that question is, what is the average spacing between the weakest false-positive sites that are scored as binding events? Third, there is good reason to have considered a wide range of TASD parameters here, not just the experimental values.

### 5 HOW TO PROBE HINDERED DIFFUSION IN THE NUCLEUS

At least four mechanisms may produce anomalous subdiffusion in the nucleus: binding, obstruction, viscoelasticity, and crowding. We consider what experiments might distinguish the first three, though a comprehensive review of hindered diffusion in the nucleus is far beyond the scope of this work.

One might be able to disentangle the obstruction effect from the others by measurements in which the tracer size is systematically varied. The basic experiments are to measure the excluded volume as a function of tracer size from superresolution images of equilibrated systems, and to measure the diffusion coefficient as a function of tracer size from SPT. These experiments would exploit the fact that geometric obstruction is very sensitive to tracer size, via the excluded volume. (The 2D case is examined in detail elsewhere (81).) The sensitivity can be pictured in terms of inverse size exclusion chromatography, in which tracers of known size are used to probe the properties of an unknown matrix (84). A complication is that in the nucleus the tracers and matrix might both be somewhat flexible. Examining obstruction in the nucleus would be a demanding experiment requiring custom preparation (or at least characterization) of a series of tracers, and a series of experiments on cells.

To isolate the effect of size, one must vary the size of the tracers while keeping other properties constant, that is, using “scalable tracers” as defined in (84). The tracer ought to be inert: no specific binding; no electrostatic interactions (no charge, minimal dipole moment); minimal van der Waals forces. The surface properties ought to be constant. The tracer ought to be spherical, not a random coil like dextran. The deformability of the tracer ought to be constant, so that variation in transport involves the deformation of the obstacles alone, not the tracers and the obstacles simultaneously. Some possibilities: Dendrimers, but they may be too small and squishy. Quantum dots would be good but even commercial ones would have to be characterized by methods of physical biochemistry. Some work has been published on this (135–138). Polystyrene spheres are highly fluorescent and available in a wide range of sizes, but nonspecific binding must be avoided – van der Waals forces might rule polystyrene spheres out. Homemade Ficoll or “Reinforced Ficoll,” with a wider range of sizes than the commercial product, would be good but would require synthesis and characterization (84). One powerful but utterly retro means of characterization in all these cases is ultracentrifugation.

In a soft system, distinguishing obstruction from binding would require comparison of the motion of the tracer with the motion of chromatin as represented by a fluorophore irreversibly bound to the chromatin. For motion of chromatin loci see for example Weber et al. (62) and Kepten et al. (65).

How might one distinguish obstruction from viscoelasticity? In the ideal case, the distinction is clear. Obstruction is by rigid obstacles rigidly attached to a rigid substrate. The interaction between tracer and obstacle is a step function, or a steep short-range potential. Diffusion is described by well-known relations from percolation theory (45, 56, 139). Viscoelasticity is described in terms of a memory function defined on the continuum, and there is a restoring force on all length scales. The distinction is less clear here. In cell biology, soft obstacles may be flexibly attached to a soft substrate. Furthermore, SPT gives high enough spatial resolution that it begins to measure the microscopic events responsible for viscoelasticity (though not, say, the conformational changes responsible for a protein spring). The shape of the potential may still be a useful distinguishing characteristic, as measured by thermal noise imaging. In this single-particle technique, the position of a tracer is measured by interferometric microscopy as the tracer moves diffusively. Inversion of the Boltzmann factor gives the potential as a function of position (140, 141).

## REFERENCES

1. Saxton, M., 1996. Anomalous diffusion due to binding: A Monte Carlo study. BIOPHYSICAL JOURNAL 70:1250–1262.

2. Saxton, M. J., 2007. A biological interpretation of transient anomalous subdiffusion. I. Qualitative model. BIOPHYSICAL JOURNAL 92:1178–1191.

3. Normanno, D., L. Boudarene, C. Dugast-Darzacq, J. Chen, C. Richter, F. Proux, O. Benichou, R. Voituriez, X. Darzacq, and M. Dahan, 2015. Probing the target search of DNA-binding proteins in mammalian cells using TetR as model searcher. NATURE COMMUNICATIONS 6:7357.

4. Caccianini, L., D. Normanno, I. Izeddin, and M. Dahan, 2015. Single molecule study of non-specific binding kinetics of Lacl in mammalian cells. FARADAY DISCUSSIONS 184:393–400.

5. Knight, S. C., L. Xie, W. Deng, B. Guglielmi, L. B. Witkowsky, L. Bosanac, E. T. Zhang, M. El Beheiry, J.-B. Masson, M. Dahan, Z. Liu, J. A. Doudna, and R. Tjian, 2015. Dynamics of CRISPR-Cas9 genome interrogation in living cells. SCIENCE 350:823–826.

6. Sternberg, S. H., S. Redding, M. Jinek, E. C. Greene, and J. A. Doudna, 2014. DNA interrogation by the CRISPR RNA-guided endonuclease Cas9. NATURE 507:62+.

7. Berg, O., and P. Von Hippel, 1985. DIFFUSION-CONTROLLED MACROMOLECULAR INTERACTIONS. ANNUAL REVIEW OF BIOPHYSICS AND BIOPHYSICAL CHEMISTRY 14:131–160.

8. Von Hippel, P., and O. Berg, 1989. FACILITATED TARGET LOCATION IN BIOLOGICAL-SYSTEMS. JOURNAL OF BIOLOGICAL CHEMISTRY 264:675–678.

9. Woringer, M., and X. Darzacq, 2018. Protein motion in the nucleus: from anomalous diffusion to weak interactions. BIOCHEMICAL SOCIETY TRANSACTIONS 46:945–956.

10. Slutsky, M., M. Kardar, and L. Mirny, 2004. Diffusion in correlated random potentials, with applications to DNA. PHYSICAL REVIEW E 69:061903.

11. Benichou, O., C. Chevalier, B. Meyer, and R. Voituriez, 2011. Facilitated Diffusion of Proteins on Chromatin. PHYSICAL REVIEW LETTERS 106:038102.

12. Imakaev, M. V., G. Fudenberg, and L. A. Mirny, 2015. Modeling chromosomes: Beyond pretty pictures. FEBS LETTERS 589:3031–3036.

13. Rosa, A., and C. Zimmer, 2014. Computational Models of Large-Scale Genome Architecture. In Hancock, R and Jeon, KW, editor, NEW MODELS OF THE CELL NUCLEUS: CROWDING, ENTROPIC FORCES, PHASE SEPARATION, AND FRACTALS, ELSEVIER ACADEMIC PRESS, SAN DIEGO, CA, volume 307 of International Review of Cell and Molecular Biology, 275–349.

14. Huet, S., C. Lavelle, H. Ranchon, P. Carrivain, J.-M. Victor, and A. Bancaud, 2014. Relevance and Limitations of Crowding, Fractal, and Polymer Models to Describe Nuclear Architecture. In Hancock, R and Jeon, KW, editor, NEW MODELS OF THE CELL NUCLEUS: CROWDING, ENTROPIC FORCES, PHASE SEPARATION, AND FRACTALS, ELSEVIER ACADEMIC PRESS, SAN DIEGO, CA, volume 307 of International Review of Cell and Molecular Biology, 443–479.

15. Amitai, A., and D. Holcman, 2017. Polymer physics of nuclear organization and function. PHYSICS REPORTS-REVIEW SECTION OF PHYSICS LETTERS 678:1–83.

16. Saxton, M., 2007. Modeling 2D and 3D Diffusion. Methods in Molecular Biology 400:295–321.

17. Saxton, M., 1994. ANOMALOUS DIFFUSION DUE TO OBSTACLES – A MONTE-CARLO STUDY. BIOPHYSICAL JOURNAL 66:394–401.

18. Forbes, C. S., M. Evans, N. A. J. Hastings, and J. B. Peacock, 2011. Statistical distributions. Hoboken, N.J: Wiley, 4th edition.

19. Newman, M., 2005. Power laws, Pareto distributions and Zipf’s law. CONTEMPORARY PHYSICS 46:323–351.

20. Saxton, M., 1993. LATERAL DIFFUSION IN AN ARCHIPELAGO – SINGLE-PARTICLE DIFFUSION. BIOPHYSICAL JOURNAL 64:1766–1780.

21. Saxton, M. J., 2008. A biological interpretation of transient anomalous subdiffusion. II. reaction kinetics. BIOPHYSICAL JOURNAL 94:760–771.

22. Rose, C., and M. D. Smith, 2002. Mathematical statistics with Mathematica. New York: Springer.

23. Bernasconi, J., H. Beyeler, and S. Strassler, 1979. ANOMALOUS FREQUENCY-DEPENDENT CONDUCTIVITY IN DISORDERED ONE-DIMENSIONAL SYSTEMS. PHYSICAL REVIEW LETTERS 42:819–822.

24. Metzler, R., J.-H. Jeon, A. G. Cherstvy, and E. Barkai, 2014. Anomalous diffusion models and their properties: non-stationarity, non-ergodicity, and ageing at the centenary of single particle tracking. PHYSICAL CHEMISTRY CHEMICAL PHYSICS 16:24128–24164.

25. Nelson, J., 1999. Continuous-time random-walk model of electron transport in nanocrystalline TiO2 electrodes. PHYSICAL REVIEWB 59:15374–15380.

26. Montroll, E., and M. Shlesinger, 1982. ON 1/F NOISE AND OTHER DISTRIBUTIONS WITH LONG TAILS (LOG-NORMAL DISTRIBUTION LEVY DISTRIBUTION PARETO DISTRIBUTION SCALE-INVARIANT PROCESS). PROCEEDINGS OF THE NATIONAL ACADEMY OF SCIENCES OF THE UNITED STATES OF AMERICA-PHYSICAL SCIENCES 79:3380–3383.

27. Mitzenmacher, M., 2004. A Brief History of Generative Models for Power Law and Lognormal Distributions. Internet Mathematics 1:226–251.

28. Malevergne, Y., V. Pisarenko, and D. Sornette, 2011. Testing the Pareto against the lognormal distributions with the uniformly most powerful unbiased test applied to the distribution of cities. PHYSICAL REVIEW E 83:036111.

29. Slutsky, M., and L. Mirny, 2004. Kinetics of protein-DNA interaction: Facilitated target location in sequence-dependent potential. BIOPHYSICAL JOURNAL 87:4021–4035.

30. Sheinman, M., O. Benichou, Y. Kafri, and R. Voituriez, 2012. Classes of fast and specific search mechanisms for proteins on DNA. REPORTS ON PROGRESS IN PHYSICS 75:026601.

31. Stewart, A. J., and J. B. Plotkin, 2012. Why Transcription Factor Binding Sites Are Ten Nucleotides Long. GENETICS 192:973+.

32. Aurell, E., A. F. d’Herouel, C. Malmnas, and M. Vergassola, 2007. Transcription factor concentrations versus binding site affinities in the yeast S-cerevisiae. PHYSICAL BIOLOGY 4:134–143.

33. Djordjevic, M., A. Sengupta, and B. Shraiman, 2003. A biophysical approach to transcription factor binding site discovery. GENOME RESEARCH 13:2381–2390.

34. Zheng, X., and J. Wang, 2015. The Universal Statistical Distributions of the Affinity, Equilibrium Constants, Kinetics and Specificity in Biomolecular Recognition. PLOS COMPUTATIONAL BIOLOGY 11:e1004212.

35. Garcia, H. G., P. Grayson, L. Han, M. Inamdar, J. Kondev, P. C. Nelson, R. Phillips, J. Widom, and P. A. Wiggins, 2007. Biological consequences of tightly bent DNA: The other life of a macromolecular celebrity. BIOPOLYMERS 85:115–130.

36. Bancaud, A., S. Huet, N. Daigle, J. Mozziconacci, J. Beaudouin, and J. Ellenberg, 2009. Molecular crowding affects diffusion and binding of nuclear proteins in heterochromatin and reveals the fractal organization of chromatin. EMBO JOURNAL 28:3785–3798.

37. Mirny, L. A., 2011. The fractal globule as a model of chromatin architecture in the cell. CHROMOSOME RESEARCH 19:37–51.

38. Loverdo, C., O. Benichou, R. Voituriez, A. Biebricher, I. Bonnet, and P. Desbiolles, 2009. Quantifying Hopping and Jumping in Facilitated Diffusion of DNA-Binding Proteins. PHYSICAL REVIEW LETTERS 102:188101.

39. Parsaeian, A., M. O. de la Cruz, and J. F. Marko, 2013. Binding-rebinding dynamics of proteins interacting nonspecifically with a long DNA molecule. PHYSICAL REVIEW E 88:040703.

40. Amitai, A., 2018. Chromatin Configuration Affects the Dynamics and Distribution of a Transiently Interacting Protein. BIOPHYSICAL JOURNAL 114:766–771.

41. Guigas, G., C. Kalla, and M. Weiss, 2007. Probing the nanoscale viscoelasticity of intracellular fluids in living cells. BIOPHYSICAL JOURNAL 93:316–323.

42. Haus, J., K. Kehr, and J. Lyklema, 1982. DIFFUSION IN A DISORDERED MEDIUM. PHYSICAL REVIEW B 25:2905–2907.

43. Bouchaud, J., and A. Georges, 1990. ANOMALOUS DIFFUSION IN DISORDERED MEDIA – STATISTICAL MECHANISMS, MODELS AND PHYSICAL APPLICATIONS. PHYSICS REPORTS-REVIEW SECTION OF PHYSICS LETTERS 195:127–293.

44. Haus, J., and K. Kehr, 1987. DIFFUSION IN REGULAR AND DISORDERED LATTICES. PHYSICS REPORTS-REVIEW SECTION OF PHYSICS LETTERS 150:263–406.

45. Klafter, J., and I. M. Sokolov, 2011. First steps in random walks: From tools to applications. Oxford: Oxford University Press.

46. Saxton, M., 1990. LATERAL DIFFUSION IN A MIXTURE OF MOBILE AND IMMOBILE PARTICLES – A MONTE-CARLO STUDY. BIOPHYSICAL JOURNAL 58:1303–1306.

47. Hinow, P., C. E. Rogers, C. E. Barbieri, J. A. Pietenpol, A. K. Kenworthy, and E. DiBenedetto, 2006. The DNA binding activity of p53 displays reaction-diffusion kinetics. BIOPHYSICAL JOURNAL 91:330–342.

48. Ellis-Davies, G. C. R., 2007. Caged compounds: photorelease technology for control of cellular chemistry and physiology. NATURE METHODS 4:619–628.

49. Lee, H.-M., D. R. Larson, and D. S. Lawrence, 2009. Illuminating the Chemistry of Life: Design, Synthesis, and Application s of “Caged” and Related Photoresponsive Compounds. ACS CHEMICAL BIOLOGY 4:409–427.

50. Jain, P. K., V. Ramanan, A. G. Schepers, N. S. Dalvie, A. Panda, H. E. Fleming, and S. N. Bhatia, 2016. Development of Light-Activated CRISPR Using Guide RNAs with Photocleavable Protectors. ANGEWANDTE CHEMIE-INTERNATIONAL EDITION 55:12440–12444.

51. Gautier, A., C. Gauron, M. Volovitch, D. Bensimon, L. Jullien, and S. Vriz, 2014. How to control proteins with light in living systems. NATURE CHEMICAL BIOLOGY 10:533–541.

52. Zhang, K., and B. Cui, 2015. Optogenetic control of intracellular signaling pathways. Trends in Biotechnology 33:92–100.

53. Kolar, K., and W. Weber, 2017. Synthetic biological approaches to optogenetically control cell signaling. CURRENT OPINION IN BIOTECHNOLOGY 47:112–119.

54. Motta-Mena, L. B., A. Reade, M. J. Mallory, S. Glantz, O. D. Weiner, K. W. Lynch, and K. H. Gardner, 2014. An optogenetic gene expression system with rapid activation and deactivation kinetics. NATURE CHEMICAL BIOLOGY 10:196+.

55. Hughes, R. M., 2018. A compendium of chemical and genetic approaches to light-regulated gene transcription. CRITICAL REVIEWS IN BIOCHEMISTRY AND MOLECULAR BIOLOGY 53:453–474.

56. Ben-Avraham, D., and S. Havlin, 2000. Diffusion and reactions in fractals and disordered systems. Cambridge: Cambridge University Press.

57. Barkai, E., Y. Garini, and R. Metzler, 2012. STRANGE KINETICS of single molecules in living cells. PHYSICS TODAY 65:29–35.

58. Hoefling, F., and T. Franosch, 2013. Anomalous transport in the crowded world of biological cells. REPORTS ON PROGRESS IN PHYSICS 76:046602.

59. Krapf, D., 2015. Mechanisms Underlying Anomalous Diffusion in the Plasma Membrane. In Kenworthy, AK, editor, LIPID DOMAINS, ELSEVIER ACADEMIC PRESS, SAN DIEGO, CA, volume 75 of Current Topics in Membranes, 167–207.

60. Sokolov, I. M., 2012. Models of anomalous diffusion in crowded environments. SOFT MATTER 8:9043–9052.

61. Mandelbrot, B., and J. Van Ness, 1968. FRACTIONAL BROWNIAN MOTIONS FRACTIONAL NOISES AND APPLICATIONS. SIAM REVIEW 10:422+.

62. Weber, S. C., J. A. Theriot, and A. J. Spakowitz, 2010. Subdiffusive motion of a polymer composed of subdiffusive monomers. PHYSICAL REVIEW E 82:011913.

63. Weiss, M., 2013. Single-particle tracking data reveal anticorrelated fractional Brownian motion in crowded fluids. PHYSICAL REVIEW E 88:010101.

64. Golan, Y., and E. Sherman, 2017. Resolving mixed mechanisms of protein subdiffusion at the T cell plasma membrane. NATURE COMMUNICATIONS 8:15851.

65. Kepten, E., I. Bronshtein, and Y. Garini, 2011. Ergodicity convergence test suggests telomere motion obeys fractional dynamics. PHYSICAL REVIEW E 83:041919.

66. Szymanski, J., and M. Weiss, 2009. Elucidating the Origin of Anomalous Diffusion in Crowded Fluids. PHYSICAL REVIEW LETTERS 103:038102.

67. Thiel, F., F. Flegel, and I. M. Sokolov, 2013. Disentangling Sources of Anomalous Diffusion. PHYSICAL REVIEW LETTERS 111:010601.

68. Izeddin, I., V. Recamier, L. Bosanac, I. I. Cisse, L. Boudarene, C. Dugast-Darzacq, F. Proux, O. Benichou, R. Voituriez, O. Bensaude, M. Dahan, and X. Darzacq, 2014. Single-molecule tracking in live cells reveals distinct target-search strategies of transcription factors in the nucleus. ELIFE 3:e02230.

69. Tafvizi, A., L. A. Mirny, and A. M. van Oijen, 2011. Dancing on DNA: Kinetic Aspects of Search Processes on DNA. CHEMPHYSCHEM 12:1481–1489.

70. Kong, M., and B. Van Houten, 2017. Rad4 recognition-at-a-distance: Physical basis of conformation-specific anomalous diffusion of DNA repair proteins. PROGRESS IN BIOPHYSICS & MOLECULAR BIOLOGY 127:93–104.

71. Normanno, D., M. Dahan, and X. Darzacq, 2012. Intra-nuclear mobility and target search mechanisms of transcription factors: A single-molecule perspective on gene expression. BIOCHIMICA ET BIOPHYSICA ACTA-GENEREGULATORY MECHANISMS 1819:482–493.

72. Bauer, M., E. S. Rasmussen, M. A. Lomholt, and R. Metzler, 2015. Real sequence effects on the search dynamics of transcription factors on DNA. SCIENTIFIC REPORTS 5:10072.

73. Veksler, A., and A. B. Kolomeisky, 2013. Speed-Selectivity Paradox in the Protein Search for Targets on DNA: Is It Real or Not? JOURNAL OF PHYSICAL CHEMISTRY B 117:12695–12701.

74. Barbi, M., C. Place, V. Popkov, and M. Salerno, 2004. Base-sequence-dependent sliding of proteins on DNA. PHYSICAL REVIEW E 70:041901.

75. Barbi, M., C. Place, V. Popkov, and M. Salerno, 2004. A model of sequence-dependent protein diffusion along DNA. JOURNAL OF BIOLOGICAL PHYSICS 30:203–226.

76. Bettridge, K. E., C. H. Bohrer, and J. Xiao, 2017. Probing the search dynamics of RNA polymerase in live E. coli cells. MOLECULAR BIOLOGY OF THE CELL 28:M202.

77. Bettridge, K., C. Bohrer, and J. Xiao, 2017. Investigating RNAP Search Dynamics in Live E. Coli Cells using Single Molecule and Statistical Methods. BIOPHYSICAL JOURNAL 112:312A.

78. Bohrer, C. H., K. Bettridge, and J. Xiao, 2017. Reduction of Confinement Error in Single-Molecule Tracking in Live Bacterial Cells Using SPICER. BIOPHYSICAL JOURNAL 112:568–574.

79. Persson, F., M. Linden, C. Unoson, and J. Elf, 2013. Extracting intracellular diffusive states and transition rates from single-molecule tracking data. NATURE METHODS 10:265–269.

80. Saxton, M., 1995. SINGLE-PARTICLE TRACKING – EFFECTS OF CORRALS. BIOPHYSICAL JOURNAL 69:389–398.

81. Saxton, M. J., 2010. Two-Dimensional Continuum Percolation Threshold for Diffusing Particles of Nonzero Radius. BIOPHYSICAL JOURNAL 99:1490–1499.

82. Chow, E., and J. Skolnick, 2017. DNA Internal Motion Likely Accelerates Protein Target Search in a Packed Nucleoid. BIOPHYSICAL JOURNAL 112:2261–2270.

83. Kang, H., Y.-G. Yoon, D. Thirumalai, and C. Hyeon, 2015. Confinement-Induced Glassy Dynamics in a Model for Chromosome Organization. PHYSICAL REVIEW LETTERS 115:198102.

84. Saxton, M. J., 2014. Wanted: Scalable Tracers for Diffusion Measurements. JOURNAL OF PHYSICAL CHEMISTRY B 118:12805–12817.

85. Cutler, P. J., M. D. Malik, S. Liu, J. M. Byars, D. S. Lidke, and K. A. Lidke, 2013. Multi-Color Quantum Dot Tracking Using a High-Speed Hyperspectral Line-Scanning Microscope. PLOS ONE 8:e64320.

86. Dross, N., C. Spriet, M. Zwerger, G. Mueller, W. Waldeck, and J. Langowski, 2009. Mapping eGFP Oligomer Mobility in Living Cell Nuclei. PLOS ONE 4:e5041.

87. Baum, M., F. Erdel, M. Wachsmuth, and K. Rippe, 2014. Retrieving the intracellular topology from multi-scale protein mobility mapping in living cells. NATURE COMMUNICATIONS 5:4494.

88. Erdel, F., M. Baum, and K. Rippe, 2015. The viscoelastic properties of chromatin and the nucleoplasm revealed by scale-dependent protein mobility. JOURNAL OF PHYSICS-CONDENSED MATTER 27:064115.

89. Brittingham, G., M. Delarue, S. Pfeffer, I. V. Surovtsev, K. J. Kennedy, S. Pinglay, I. Gutierrez, D. Sang, G. Poterewicz, J. Chung, J. T. Groves, C. Jacobs-Wagner, B. D. Engel, and L. J. Holt, 2017. mTORC1 controls rheology and phase separation by tuning ribosome concentration. MOLECULAR BIOLOGY OF THE CELL 28:P2593.

90. Pinglay, S., G. Brittingham, M. Delarue, K. J. Kennedy, G. Poterewicz, and L. J. Holt, 2017. Genetically Encoded Multimeric nanoparticles (GEMs) to visualize the biophysical properties of the nucleus. MOLECULAR BIOLOGY OF THE CELL 28:P3116.

91. Delarue, M., G. P. Brittingham, S. Pfeffer, I. V. Surovtsev, S. Pinglay, K. J. Kennedy, M. Schaffer, J. I. Gutierrez, D. Sang, G. Poterewicz, J. K. Chung, J. M. Plitzko, J. T. Groves, C. Jacobs-Wagner, B. D. Engel, and L. J. Holt, 2018. mTORC1 Controls Phase Separation and the Biophysical Properties of the Cytoplasm by Tuning Crowding. CELL 174:338+.

92. Giessen, T. W., 2016. Encapsulins: microbial nanocompartments with applications in biomedicine, nanobiotechnology and materials science. CURRENT OPINION IN CHEMICAL BIOLOGY 34:1–10.

93. King, N. P., J. B. Bale, W. Sheffler, D. E. McNamara, S. Gonen, T. Gonen, T. O. Yeates, and D. Baker, 2014. Accurate design of co-assembling multi-component protein nanomaterials. NATURE 510:103+.

94. Qian, L., and E. Kussell, 2016. Genome-Wide Motif Statistics are Shaped by DNA Binding Proteins over Evolutionary Time Scales. PHYSICAL REVIEW X 6:041009.

95. Buchanan, M., 2016. Focus: Evolution Thins Out Distracting DNA. 30 Jul 2019. https://physics.aps.org/articles/v9/119.

96. Elf, J., G.-W. Li, and X. S. Xie, 2007. Probing transcription factor dynamics at the single-molecule level in a living cell. SCIENCE 316:1191–1194.

97. Elf, J., and I. Barkefors, 2019. Single-Molecule Kinetics in Living Cells. Annual Review of Biochemistry 88:000–000.

98. Pederson, T., 2014. The Nuclear Physique. In Hancock, R and Jeon, KW, editor, NEW MODELS OF THE CELL NUCLEUS: CROWDING, ENTROPIC FORCES, PHASE SEPARATION, AND FRACTALS, ELSEVIER ACADEMIC PRESS, SAN DIEGO, CA, volume 307 of International Review of Cell and Molecular Biology, 1–13.

99. Gorman, J., F. Wang, S. Redding, A. J. Plys, T. Fazio, S. Wind, E. E. Alani, and E. C. Greene, 2012. Single-molecule imaging reveals target-search mechanisms during DNA mismatch repair. PROCEEDINGS OF THE NATIONAL ACADEMY OF SCIENCES OF THE UNITED STATES OF AMERICA 109:E3074–E3083.

100. Hacker, W. C., S. Li, and A. H. Elcock, 2017. Features of genomic organization in a nucleotide-resolution molecular model of the Escherichia coli chromosome. NUCLEIC ACIDS RESEARCH 45:7541–7554.

101. Tanay, A., 2006. Extensive low-affinity transcriptional interactions in the yeast genome. GENOME RESEARCH 16:962–972.

102. Hahn, M., J. Stajich, and G. Wray, 2003. The effects of selection against spurious transcription factor binding sites. MOLECULAR BIOLOGY AND EVOLUTION 20:901–906.

103. Kirchheim, R., 1988. HYDROGEN SOLUBILITY AND DIFFUSIVITY IN DEFECTIVE AND AMORPHOUS METALS. PROGRESS IN MATERIALS SCIENCE 32:261–325.

104. Cameron, L., and C. Sholl, 1999. The average jump rate and diffusion in disordered systems. JOURNAL OF PHYSICS-CONDENSED MATTER 11:4491–4497.

105. Dvoyashkin, M., A. Khokhlov, S. Naumov, and R. Valiullin, 2009. Pulsed field gradient NMR study of surface diffusion in mesoporous adsorbents. MICROPOROUS AND MESOPOROUS MATERIALS 125:58–62.

106. Berg, O., and P. Von Hippel, 1987. SELECTION OF DNA-BINDING SITES BY REGULATORY PROTEINS – STATISTICAL-MECHANICAL THEORY AND APPLICATION TO OPERATORS AND PROMOTERS. JOURNAL OF MOLECULAR BIOLOGY 193:723–743.

107. Roider, H. G., A. Kanhere, T. Manke, and M. Vingron, 2007. Predicting transcription factor affinities to DNA from a biophysical model. BIOINFORMATICS 23:134–141.

108. Ruan, S., and G. D. Stormo, 2017. Inherent limitations of probabilistic models for protein-DNA binding specificity. PLOS COMPUTATIONAL BIOLOGY 13:e1005638.

109. Wasson, T., and A. J. Hartemink, 2009. An ensemble model of competitive multi-factor binding of the genome. GENOME RESEARCH 19:2101–2112.

110. Saxton, M., 2001. Anomalous subdiffusion in fluorescence photobleaching recovery: A Monte Carlo study. BIOPHYSICAL JOURNAL 81:2226–2240.

111. Bulnes, F., V. Pereyra, and J. Riccardo, 1998. Collective surface diffusion: n-fold way kinetic Monte Carlo simulation. PHYSICAL REVIEW E 58:86–92.

112. Press, W. H., S. A. Teukolsky, W. T. Vetterling, and B. P. Flannery, 1992. Numerical Recipes in FORTRAN. The Art of Scientific Computing. Cambridge: Cambridge University Press, 2nd edition.

113. Ellery, A. J., R. E. Baker, and M. J. Simpson, 2016. Communication: Distinguishing between short-time non-Fickian diffusion and long-time Fickian diffusion for a random walk on a crowded lattice. JOURNAL OF CHEMICAL PHYSICS 144:171104.

114. Wolfram Language & System Documentation Center, 2019. Random Number Generation. 30 Jul 2019. https://reference.wolfram.com/language/tutorial/RandomNumberGeneration.html.

115. Wolfram, S., 2016. Idea makers: Personal perspectives on the lives & ideas of some notable people. Champaign, Illinois: Wolfram Media, Inc.

116. Mathematica Stack Exchange, 2012. Quality of random numbers. 30 Jul 2019. https://mathematica.stackexchange.com/questions/3208/quality-of-random-numbers.

117. Wikipedia, 2019. Generalized mean. 30 Jul 2019. https://en.wikipedia.org/wiki/Generalized_mean.

118. Cantrell, D. W., and E. W. Weisstein. Power mean. 30 Jul 2019. MathWorld–A Wolfram Web Resource. http://mathworld.wolfram.com/PowerMean.html.

119. Zaninetti, L., 2017. A left and right truncated lognormal distribution for the stars. Advances Astrophys 2:197–213.

120. Pottier, N., 2011. Relaxation time distributions for an anomalously diffusing particle. PHYSICA A-STATISTICAL MECHANICS AND ITS APPLICATIONS 390:2863–2879.

121. Liebovitch, L., and T. Toth, 1991. DISTRIBUTIONS OF ACTIVATION-ENERGY BARRIERS THAT PRODUCE STRETCHED EXPONENTIAL PROBABILITY-DISTRIBUTIONS FOR THE TIME SPENT IN EACH STATE OF THE 2 STATE REACTION A-REVERSIBLE-B. BULLETIN OF MATHEMATICAL BIOLOGY 53:443–455.

122. Metzler, R., J. Klafter, J. Jortner, and M. Volk, 1998. Multiple time scales for dispersive kinetics in early events of peptide folding. CHEMICAL PHYSICS LETTERS 293:477–484.

123. Ostrowsky, N., D. Sornette, P. Parker, and E. Pike, 1981. EXPONENTIAL SAMPLING METHOD FOR LIGHT-SCATTERING POLYDISPERSITY ANALYSIS. OPTICA ACTA 28:1059–1070.

124. Berberan-Santos, M., E. Bodunov, and B. Valeur, 2005. Mathematical functions for the analysis of luminescence decays with underlying distributions 1. Kohlrausch decay function (stretched exponential). CHEMICAL PHYSICS 315:171–182.

125. Barone, P., A. Ramponi, and G. Sebastiani, 2001. On the numerical inversion of the Laplace transform for nuclear magnetic resonance relaxometry. INVERSE PROBLEMS 17:77–94.

126. Plonka, A., J. Kroh, and Y. Berlin, 1988. PHOTODISSOCIATION OF CARBON MONOXY MYOGLOBIN – KINETICS OF CARBON-MONOXIDE REBINDING. CHEMICAL PHYSICS LETTERS 153:433–435.

127. Istratov, A., and O. Vyvenko, 1999. Exponential analysis in physical phenomena. REVIEW OF SCIENTIFIC INSTRUMENTS 70:1233–1257.

128. Epstein, C. L., and J. Schotland, 2008. The bad truth about Laplace’s transform. SIAM REVIEW 50:504–520.

129. Lee, S., 2003. Correction-to-scaling of random walks in disordered media. INTERNATIONAL JOURNAL OF MODERN PHYSICS B 17:4867–4881.

130. Shlesinger, M., 1988. FRACTAL TIME IN CONDENSED MATTER. ANNUAL REVIEW OF PHYSICAL CHEMISTRY 39:269–290.

131. Berg, O., R. Winter, and P. Von Hippel, 1981. DIFFUSION-DRIVEN MECHANISMS OF PROTEIN TRANSLOCATION ON NUCLEIC-ACIDS.1. MODELS AND THEORY. BIOCHEMISTRY 20:6929–6948.

132. Halford, S., and J. Marko, 2004. How do site-specific DNA-binding proteins find their targets? NUCLEIC ACIDS RESEARCH 32:3040–3052.

133. Nemirovsky, A., H. Martin, and M. Coutino-Filho, 1990. UNIVERSALITY IN THE LATTICE-COVERING TIME PROBLEM. PHYSICAL REVIEW A 41:761–767.

134. Brummelhuis, M., and H. Hilhorst, 1991. COVERING OF A FINITE LATTICE BY A RANDOM-WALK. PHYSICA A-STATISTICAL MECHANICS AND ITS APPLICATIONS 176:387–408.

135. Howarth, M., W. Liu, S. Puthenveetil, Y. Zheng, L. F. Marshall, M. M. Schmidt, K. D. Wittrup, M. G. Bawendi, and A. Y. Ting, 2008. Monovalent, reduced-size quantum dots for imaging receptors on living cells. NATURE METHODS 5:397–399.

136. Lees, E. E., M. J. Gunzburg, T.-L. Nguyen, G. J. Howlett, J. Rothacker, E. C. Nice, A. H. A. Clayton, and P. Mulvaney, 2008. Experimental determination of quantum dot size distributions, ligand packing densities, and bioconjugation using analytical ultracentrifugation. NANO LETTERS 8:2883–2890.

137. Sperling, R. A., T. Pellegrino, J. K. Li, W. H. Chang, and W. J. Parak, 2006. Electrophoretic separation of nanoparticles with a discrete number of functional groups. ADVANCED FUNCTIONAL MATERIALS 16:943–948.

138. Lin, C.-A. J., R. A. Sperling, J. K. Li, T.-Y. Yang, P.-Y. Li, M. Zanella, W. H. Chang, and W. J. Parak, 2008. Design of an amphiphilic polymer for nanoparticle coating and functionalization. SMALL 4:334–341.

139. Hughes, B. D., 1996. Random walks and random environments. Volume 2. Random environments. Oxford: Clarendon Press.

140. Tischer, C., S. Altmann, S. Fisinger, J. Horber, E. Stelzer, and E. Florin, 2001. Three-dimensional thermal noise imaging. APPLIED PHYSICS LETTERS 79:3878–3880.

141. Bartsch, T. F., M. D. Kochanczyk, E. N. Lissek, J. R. Lange, and E.-L. Florin, 2016. Nanoscopic imaging of thick heterogeneous soft-matter structures in aqueous solution. NATURE COMMUNICATIONS 7:12729.

